# Single particle combinatorial multiplexed liposome fusion mediated by DNA

**DOI:** 10.1101/2021.01.19.427313

**Authors:** Mette Galsgaard Malle, Philipp M. G. Löffler, Søren S.-R. Bohr, Magnus Berg Sletfjerding, Nikolaj Alexander Risgaard, Simon Bo Jensen, Min Zhang, Per Hedegaard, Stefan Vogel, Nikos S. Hatzakis

## Abstract

Combinatorial high throughput methodologies are central for both screening and discovery in synthetic biochemistry and biomedical sciences. They are, however, often reliant on large scale analyses and thus limited by long running time and excessive materials cost. We herein present **S**ingle **PAR**ticle **C**ombinatorial multiplexed **L**iposome fusion mediated by **D**NA (SPARCLD), for the parallelized, multi-step and non-deterministic fusion of individual zeptoliter nanocontainers. We observed directly the efficient (>93%), and leakage free stochastic fusion sequences for arrays of surface tethered *target* liposomes with six freely diffusing populations of *cargo* liposomes, each functionalized with individual lipidated ssDNA (LiNA) and fluorescent barcoded by distinct ratio of chromophores. The stochastic fusion results in distinct permutation of fusion sequences for each autonomous nanocontainer. Real-time TIRF imaging allowed the direct observation of >16000 fusions and 566 distinct fusion sequences accurately classified using machine learning. The high-density arrays of surface tethered target nanocontainers ∼42,000 containers per mm^2^ offers entire combinatorial multiplex screens using only picograms of material.

## Introduction

High-throughput (HTP) combinatorial methodologies are essential for accelerating synthetic biochemical technologies and discovery platforms, to reduce the time and expenses for studies with large parameter space and in-depth analysis. Their use relies primarily on microarrays^1^, lab-on-a-chip systems^2^, microfluidics^3^, parallel pipetting^4^, or robotic assisted methodologies^5^ which greatly minimize manpower and offers automated parallelized screening (10^3^–10^6^) of small molecules, albeit requiring large quantities of material and considerable running time. To reduce the cost and material, Ultra-Miniaturized assays have been developed. They may involve picoliter lipid droplets screening of metagenomic libraries using microfluidics^3^, or parallelized subattoliter content mixing via membrane fusion^6^. Efficient membrane fusion have previously been accomplished using both reconstituted SNARE proteins^7^, charged lipids^6,8^, SNARE-mimics such as lipopeptides^9,10^ or, recently, lipidated DNA (LiNA) oligomers^11,12^. The facile programmability and exchangeability of DNA sequences^13–16^ combined with DNA mediated membrane fusion can offer multi-step content mixing^11^ and the possibility of combining fusion chemical cascades using fluorescent microscopy techniques, but to date they are primarily used for sequential mixing.

Fluorescence microscopy is a powerful and sensitive detection tool^17–19^, but its capability for multiplexing is limited by the spectral overlap between chromophores, restricting quantitative imaging to a handful of fluorescent colors^20,21^. Fluorescent barcoding technologies can overcome this, offering some success in multiplexing for both *in vitro* and *in vivo* imaging. *In vitro* barcodes such as quantum dot-based microbeads relying on intensity encoding offer the potential of a massive color pallet, but are limited by their large size and need for functionalization ^22,23^. Spectral encoding using stimulated Raman scattering techniques can provide a pallet of 30 distinct frequencies^24,25^, however their reliance on highly sensitive detection reduces their capacity for imaging dynamic processes. *In vivo* simultaneous monitoring of multiple compartments in cells has been achieved by a spatial encoding relying on super resolution imaging together with combinatorial labelling of mRNAs^26^, however they are reliant on massive randomized labeling and require sensitive imaging. Combinatorial expression of fluorescent proteins^27^, may on the other hand reach up to 90 colors, albeit phototoxicity and their dependence on sensitive imaging may limit their applicability. Nano-scale geometric barcodes^15^ are a novel strategy for multiplexed labelling of multiple molecular species both *in vivo* and *in vitro*, but they need super-resolution imaging challenging their use in dynamic systems.

Here we present **S**ingle **PAR**ticle **C**ombinatorial **L**iposome fusion mediated by **D**NA (SPARCLD) for multiplexed cargo delivery of attoliter lipidic nanocontainers. The method combines chromophore ratio labelling creating distinct identity encoding barcodes for each DNA encoded nanocontainer and the use of complementary LiNA mediated nanocontainer fusion. Using TIRF microscopy allowed the parallelized imaging of ∼8800 individual target lipidic containers^17–19,28,29^ undergoing >16,000 fusion events with barcoded cargo nanocontainers and up to seven successive rounds of fusion. The fusion sequence is completely stochastic allowing >550 distinct permutations that are directly recorded and precisely classified by machine learning analysis. The assay dimensions allow approximately 42,000 target containers per square millimeter of microscope surface, and thus highly parallel recording results in thousands of nanocontainer experiments within an hour. SPARCLD transforms stochasticity from a prohibitive problem in conventional assays into an experimental advantage and an enabling technology for multiplexing offering the direct high throughput screening or building of synthetic biopolymers, such as carbohydrates and nucleic acids or for drug screening or epitope mapping, reducing both reagents and time.

## Results

To attain the multiplexed combinatorial fusion we combined the single stranded lipidated DNA (LiNA) functionalization technology^11^ with single liposome fluorescent readout^17,30,19^. We produced arrays of *target* nanocontainers by tethering liposomes to a passivated microscope surface using a neutravidin/biotin protocol^30^ (see micrograph Fig. 1a). This methodology maintains the spherical topology of liposomes, their low membrane permeability during immobilization and enables their unhindered interaction with biomolecules^17,31^. Each target liposome was membrane-labeled using 3,3’-dioctadecyloxacarbocyanine perchlorate (DiO) and functionalized with six different LiNA sequences (**A** to **F**). Using a total internal reflection (TIRF) microscope, hundreds of target liposomes per field of view were recorded in parallel (Fig. 1a), while extracting their sub-resolution dimensions, volumes as well as their spatial localization with nanometer precision^30^.

**Figure 1:**
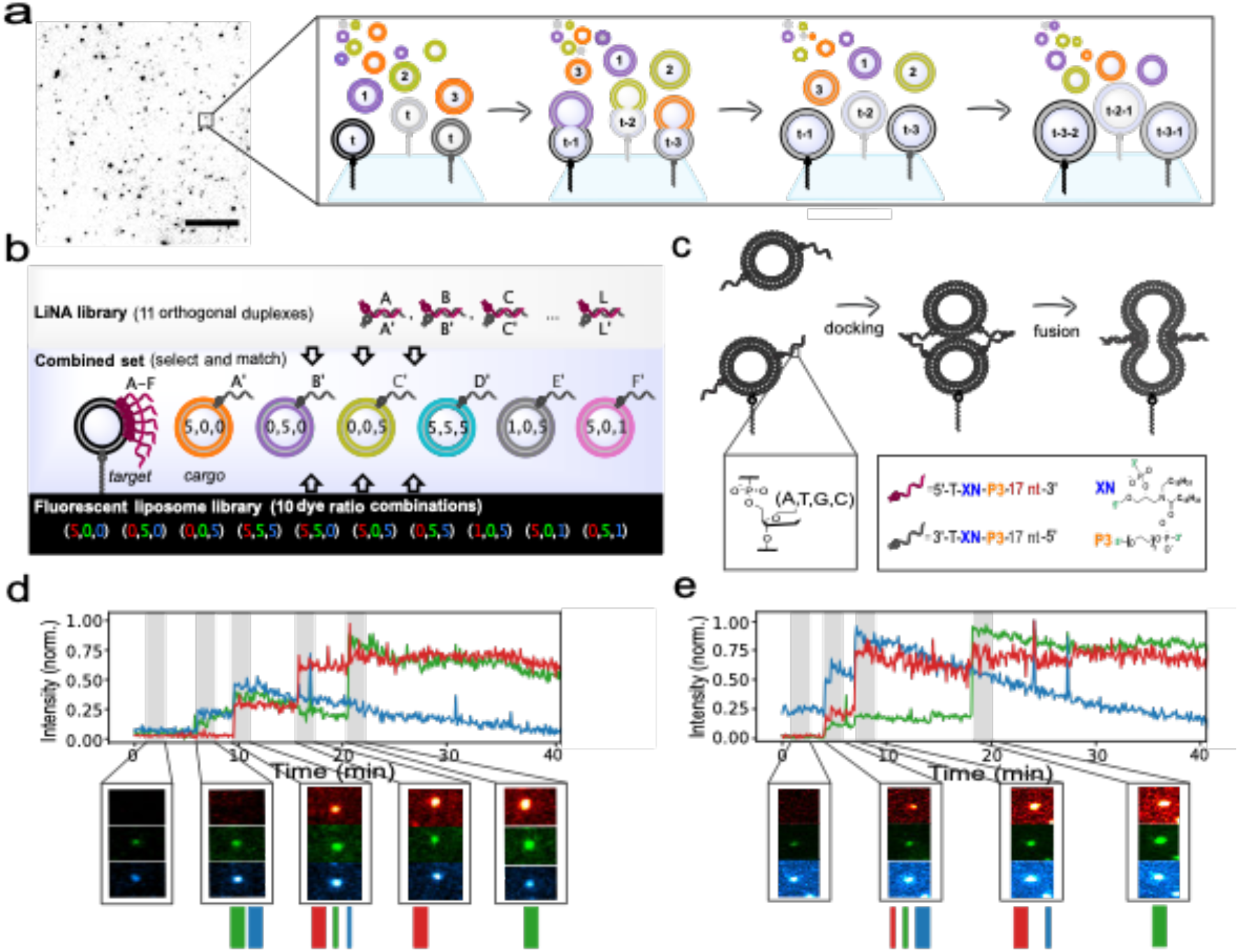
Combinatorial Liposome fusion mediated by DNA for the parallelized, fusion with stochastic sequence of individual zeptoliter lipid nanocontainers. (a) Typical micrograph of target liposomes tethered to a PLL-PEG passivated surface. Varying intensities originate from the polydisperse size distribution of the liposomes. A zoom in cartoon representation illustrates stochastic fusion events monitored on the single-particle level that can be used to detect thousands of individual DNA programmable liposome fusion events in a stochastic and multiplexed manner using TIRF microscopy and automated data analysis. *Scale bar is 10 µm*. (b) Target liposomes, each loaded with six lipidated ssDNA sequences (LiNA’s) are immobilized on the surface. Freely diffusing cargo liposomes, each functionalized with one of the six complementary LiNAs, were barcoded with a distinct ratio of up to three types of fluorescently labeled lipids. This resulted in six distinct barcodes denoted as relative Red-Green-Blue ratios, that are easily expandable to 10 barcodes and distinct up to 11 complementary pairs of LiNA sequences (see Supplementary Fig. 6 and 7). (c) The complementary LiNA sequences between target/cargo are designed to facilitate fusion of membranes by a zipper-like hybridization forcing close proximity. (d-e) Representative single particle time traces and the corresponding snapshots of a series of raw microscope images displaying two otherwise identical target liposomes undergoing repetitive fusion, within a single field of view. Data highlight the stochasticity of cargo identity, sequence and the number of repetitive fusions. Trace (d) shows four repetitive fusions, while trace (e) shows three. Precise target identity, shown as the barcodes below the snapshots, was attained by three channel signal integration and machine learning classification (see Supplementary Fig. 5 for additional fusion traces).

In order to achieve the combinatorial fusion of *cargo* nanocontainers to target nanocontainers we prepared six different populations of cargo liposomes each loaded with one out of six complementary LiNA counterparts (**A’** to **F’**) to the target liposomes. Identity encoding for each of the six cargo populations was reached by ‘intensity barcoding’ using a distinct ratio of up to three fluorescently labeled lipids (Red; R, ATTO-655-DOPE, ‘green’; G, ATTO-550-DOPE and ‘blue’; B, DiO), creating distinct spectral signatures (Fig 1b). The complementary LiNA strands would hybridize in a zipper-like design bringing the bilayers into contact and facilitating efficient fusion of the membrane (Fig. 1c), as we and others have recently shown^12,32^ (see Supplementary Fig. 1). We designed single stranded LiNA sequences to hybridize selectively into orthogonal duplexes with minimal crosstalk (see Methods and supplementary figure 2). Surface passivation of the microscope glass surface minimized the non-specific binding of cargos to the surface (see Supplementary Fig. 3). Parallel three-color imaging allowed the direct real-time monitoring of each cargo liposome docking and delivering its content to each target liposome.

In a typical experiment, a mixture containing an equal amount of the six types of cargo liposomes was added into the microscope chamber slide containing surface tethered target liposomes (see Fig 1a). Cargo liposomes will either i) transiently dock with liposomes for one to two frames defined as kiss-and-run events (see spikes in Fig. 1d-e) or ii) irreversibly dock for prolonged time which will lead to fusion (see step-like signal increases, Fig. 1d-e). Control experiments with non-complementary LiNA between cargo and target liposomes showed minimal (4.8±0.9% of targets) irreversible docking events (Supplementary Fig. 4). Each target liposome can display multiple successive fusion events with any of the six barcoded cargo vesicles. Figures 1d and 1e show typical time traces on two neighboring, and otherwise identical, target liposomes, exhibiting four and three fusion events, respectively (see Supplementary Fig. 5 for more examples). For each fusion event, the step-like signal increases in the respective detection channel(s) correspond to the spectral signature of the cargo liposome and is assigned to the cargo population along with the LiNA it carries, via the intensity barcoding. Consequently, the sequence of fusion events for the entire time trace of each target liposome can be reconstructed. Because each target liposome constitutes an autonomous experiment and the fusion sequence is stochastic, neighboring target liposomes can have a completely different sequence of repetitive fusion events (see Fig. 1) that can be recorded in parallel and in real time using a TIRF microscope.

### Precise and automated classification of several successive events by machine learning

To recognize and classify each individual fused cargo liposome based on the fluorescent barcode we trained and applied a supervised machine learning (ML) algorithm based on extreme gradient boosted decision tree (see methods). Each fluorescent barcode contains specific labelling ratios of lipidated fluorophores generating an RGB coded signal in three microscope channels of ‘red’ (R), ‘green’ (G) and ‘blue’ (B) generating the identity encoding. Six fluorescent barcode populations were selected for optimal recognition and method construction and demonstration (see supplementary figure 6 for full intensity library and supplementary figure 7 for prediction accuracy and Supplementary Fig. 8 for barcode selection). The model was re-trained on a total of 44,000 liposomes from individual imaging of the six selected barcodes (see Fig 2a) and then used to classify each individual cargo liposome during experiments where all six cargo populations were available for docking and fusion.

**Figure 2:**
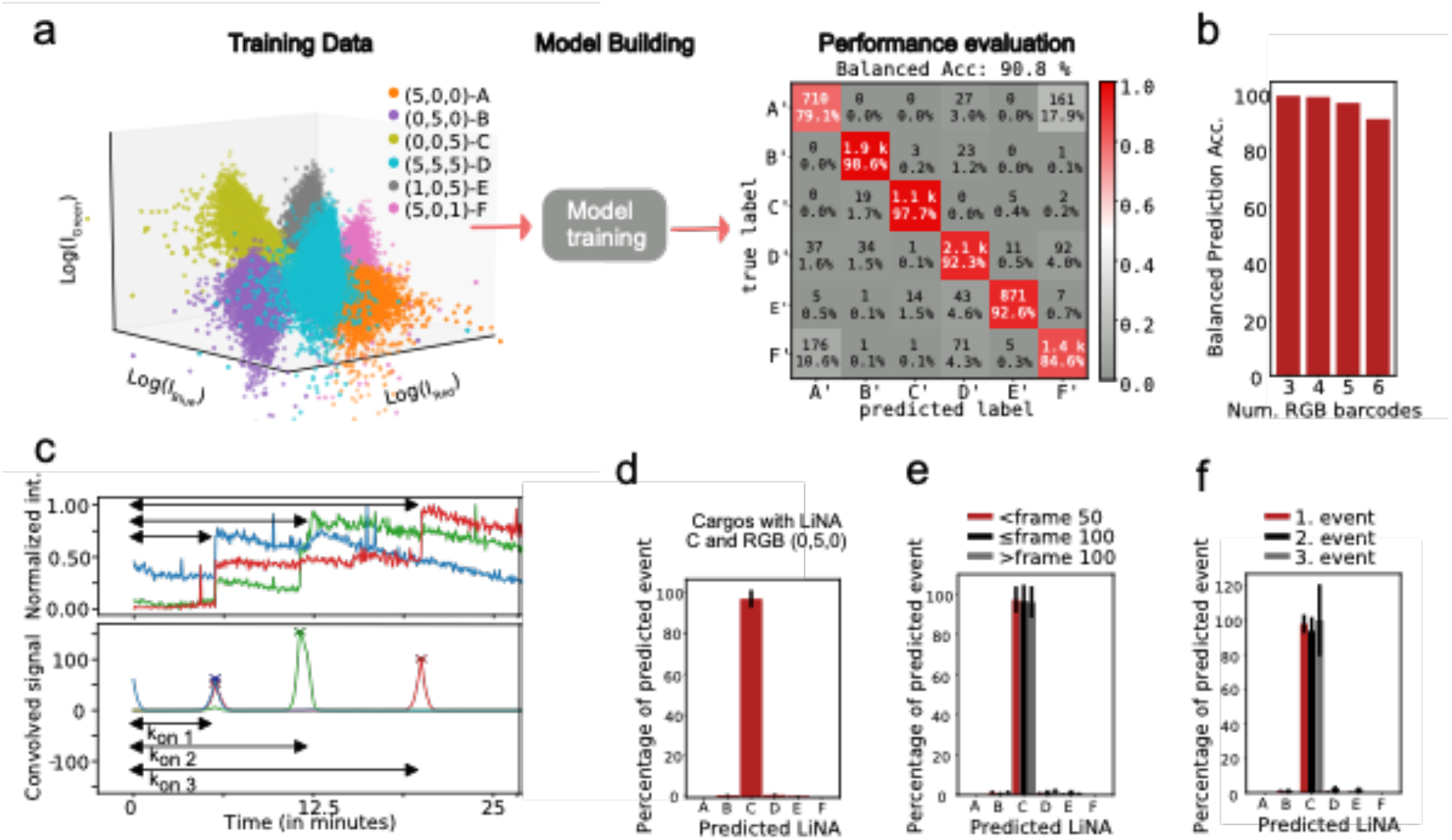
Classification Accuracy of barcoded liposomes using supervised Machine learning. (a) 3D plot of intensities in the three channels for the six barcoded liposome populations used for ML training (see Supplementary Fig. 8 for selection from the initial superset with ten barcodes). Each of the six populations contains a specific ratio of Red, Green, and Blue lipid-conjugated chromophores (R,G,B). The intensities in each channel were used for supervised model training using an extreme gradient boosted tree (N=44,000). Evaluation for the ML model is shown in the confusion matrix, displaying the classification accuracy for each of the six barcode populations. A balanced prediction accuracy of 90.8% was reached. (b) Balanced accuracy for classification for three to six barcoded liposome populations. Classification relies on supervised models and was also trained for subset of the recorded dataset (see Supplementary Fig. 9 for subset confusion matrices). c) Raw single particle trajectories of three successive liposome fusion events. If successful fusion events were registered (signal persisting more than 30 frames or 3 minutes) the barcode of the incoming cargo vesicle was classified (see Methods). The number of successive events as well the respective waiting times (k_on_) between them are extracted for thermodynamic characterization. (d) Experimental validation of the classification model using one cargo liposome barcode (LiNA C and barcode (0,0,5) as ground truth. The barcode was classified correctly 96.7±4.3 % of the time (See Supplementary Fig. 12 for further tests using LiNA D and (5,5,5) barcoded liposomes). (e) The classification accuracy remained practically identical independent on fusion occurring before frame 50, between 50 and 100 or after 100, ruling out bleaching as a potential issue for the classification model (f) The classification accuracy was independent of the number of successive fusion events (note the larger error bar for the third successive event, as it is based on fewer events). Intensity variations due to multi-color fusion and signal crosstalk did not significantly affect the accuracy of the classification method.

We evaluated the accuracy of the supervised classification model using the confusion matrix in Fig. 2a. Each row represents the predicted classification of events and each column the ground truth data. The diagonals display the number of correctly classified barcodes and the accuracy, while off diagonal features represent the misclassified barcodes. The classification accuracy exceeded 90 % in most cases with minor misclassification for liposomes with LiNA A’ and barcode (5,0,0) and some liposomes with LiNA F’ and barcode (5,0,1). The balanced accuracy for this trained model, using extreme gradient boosted trees is 90.8 %. Expectedly, the classification accuracy depends on the number of classes, and is improved for fewer barcode classes, as seen in Fig. 2b (See Supplementary Fig. 9 for confusion matrices for all the subpopulations).

Docking event detection and classification was performed using digital signal convolution. The method will exclusively pick up docking for prolonged time, and not kiss-and-run events due to the small integral for a single frame event versus step-function change (see Methods and supplementary Fig. 10). The background corrected and integrated raw signal of each detected peak in all three channels was used for barcode classification by the ML model. The method provides information on the order of cargo fusion events, the dwell time between successive events as well as the nanoscale dimensions of both cargo and target nanocontainers.

Several controls with ground truth data confirmed the precise event detection and classification using our ML model. The predicted accuracy of ML model to classify LiNA-C’ docking (RGB barcode 0,5,0) is 97.7% (see Fig. 2a). To experimental validate this, we recorded directly the fusion of LiNA-C to target liposomes engrafted with all LiNA. Fusion events were correctly assigned in 96.7±4.3 % of events, confirming the model’s high classification accuracy (see Fig. 2d). Liposome preparation, day-to-day variations in microscope/optics alignment and imaging conditions introduced no measurement bias in classification (see Supplementary Fig. 11). We found identical classification accuracy independently of whether fusion occurs in the first (<50 frames) or last part (>100 frames) of the experiments (Fig. 2e), supporting that chromophore bleaching does not bias the ML model. The classification accuracy was also found to be robust in case of repeated fusion events (see Fig. 2f) and independent of the type of barcoded liposomes (see Supplementary Fig. 12 for data (5,5,5) and LiNA D’). In summary the classification model precisely reported the RGB barcoded identity invariantly of potential bleaching, arrival time and number of successive events of the barcoding cargo liposomes, emphasizing it as a robust, rapid and reproducible classification.

### Quantitative fusion and leakage free delivery of cargo for content mixing

A quantitative, leakage-free fusion is crucial for any combinatorial multi-step cargo delivery assay. To measure this, we loaded target liposomes with the enzyme β-glucosidase (βGlu, *Aspergillus Niger*) and cargo liposomes with the pro-fluorescent substrate, Fluorescein Di-β-D-Glucopyranoside (FDGlu, see figure 3a). Successful fusion delivers the FDGlu cargo via content mixing and triggers an enzymatic reaction that hydrolyzes FDGlu to fluorescein (see supplementary figure 13-16 for bulk experiment controls). The resulting fluorescence increase was accurately detected by the sensitive microscopy setup. Labeling target liposomes with ATTO-550-DOPE and cargo liposomes with ATTO-655-DOPE allowed the synchronous recording of both cargo liposomes docking, and content mixing: Docking results in a clear single step increase in the red channel and fluorescein production upon fusion in an increase in the blue channel (see Fig. 3b, representative trace in Fig. 3c, and Supplementary Fig. 17-19 for additional traces). Fusion occurred faster than the temporal resolution^6,33,34^ (21 seconds per cycle) resulting in a one-step product signal increase (see fig. 3c) and was present for all target and cargo liposomes sizes (∼30-300nm) (see Supplementary Fig. 20). Interestingly, the docking and fusion kinetics depended on the LiNA sequence (see Supplementary Fig. 21). Successful fusion and specific LiNA mediated cargo delivery were confirmed by a fluorometric assay in bulk solution (see Supplementary Fig. 22). Single particle readouts allowed the deconvolution of the individual docking and fusion events that are averaged out in conventional measurements due to ensemble averaging of large numbers of concurrent events.

**Figure 3:**
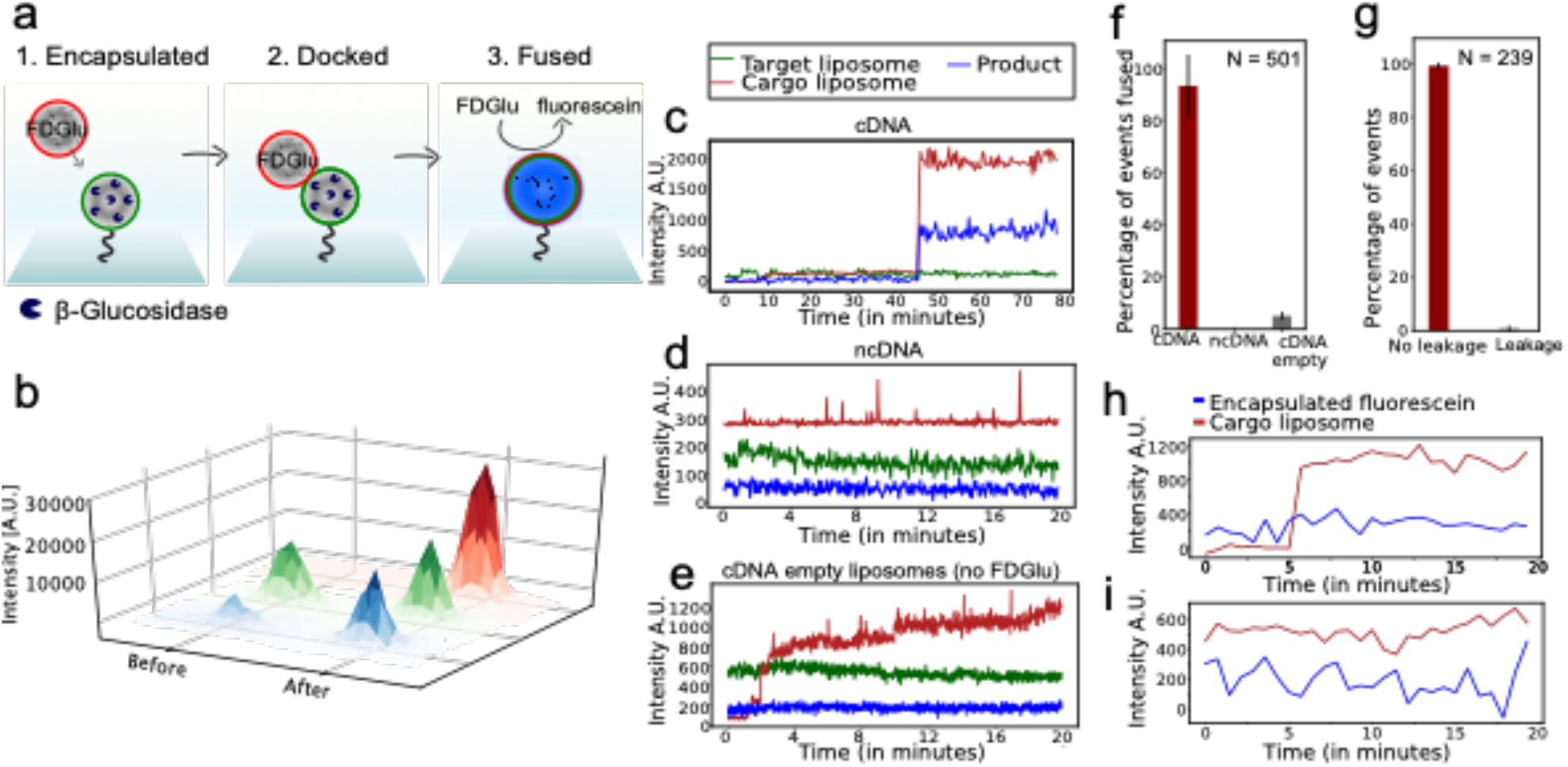
Quantitative and specific content mixing of subattolitre lipid nanocontainers. (a) Schematic illustration of real time measurements of quantitative content mixing at the individual liposome level. Target liposomes are loaded with β-glucosidase and labeled with ATTO-550-lipid for localization. Cargo liposomes are labeled with ATTO-655-lipid and loaded with a pro-fluorescent substrate, FDGlu. (b) 3D visualizations of raw snapshots of zoomed microscope images for a single liposome in all three channels prior to docking and after docking/fusion. The target liposome is localized by the green channel. Cargo docking will result in a single step increase in the red channel. Successful fusion will trigger delivery of substrate to the enzyme and thereby formation of the fluorescent product fluorescein, causing an increase in the blue channel. Product formation will rapidly occur in time frames below the temporal resolution, as seen from the instant increase in the blue channel. (c-e) Representative time traces of single target liposomes. (c) Cargo liposomes containing LiNA complementary to the target LiNAs. Cargo docking (red) results in fusion, content mixing and product formation (blue). (d) Cargo liposomes containing LiNA non-complementary to the target LiNAs showing several kiss-and-run events without successful fusion. (e) Unloaded cargo liposomes containing complementary LiNA results in docking and fusion without any product formation due to lack of substrate. (see Supplementary Fig. 17-19 for additional traces). (f) Quantification of docking and fusion efficiency: 93.2 ± 12.0 % of docked liposomes with complementary LiNA undergo fusion but 0% undergo fusion for non-complementary encoding. Cargo liposomes without FDGlu but with complementary LiNA encoding showed 5.1±1.5%, serving as false-positive control, N=501 (g) Quantification of leakage: 99.23% of the cases upon fusion with a cargo liposome are leakage free, N=239) (h) A representative leakage assay time trace, showing a single docking event (red) but no leakage of the encapsulated fluorescein in the target liposome (blue) (i) A representative time trace of a target liposome (blue) not subject to fusion showing no leakage. Both traces are recorded using sensitive low temporal resolution to avoid false positives due to product bleaching.

Quantitative analysis revealed 91.6±4.2 % of the imaged target liposomes to undergo one or more specific docking event with cargo liposomes carrying complementary LiNAs. 93.2 ± 12.0 % (N=501) of the docked liposomes successfully fused and initiated the enzymatic reaction at biologically relevant temperatures (37 °C) (see Fig. 3f). Interestingly, we found that 77.1 ± 8.8% of liposomes (N=11791) successfully encapsulated the enzyme^30^ (see Supplementary Fig. 23 and Methods). In ensemble assays, where the effects of encapsulation efficiency, docking efficiency and fusion efficiency are convoluted, the apparent fusion efficiency would be lower^11,35^. The practically quantitative progression from docking to fusion confirmed that successive docking events using the SPARCLD fusion methodology will lead to successful cargo delivery in each nanoreactor.

Several control experiments confirmed the specificity of LiNA mediate fusion. Non-specific docking of cargo to target engrafted with non-complementary LiNA (LiNA-D on both), was only 1.9±0.8%. The transient spikes (1-2 frames) (Fig. 3d) observed correspond to kiss-and-run events. Empty cargo liposomes without encapsulated FDGlu substrate but otherwise identical amounts of complementary LiNA, showed similar high docking efficiency with complimentary targets (90.0±6.1%) and only 5.1±1.5% displayed a measurable increase in product channel. This low ratio of false positive events may originate primarily from bleed-through from docked cargo liposomes, as three channels are imaged synchronously, and the high-sensitivity setup needed for detecting enzymatic product (see Fig. 3e and 3f). None of the transiently docked vesicles in the control experiment resulted in fusion, underlining the high fidelity of DNA-mediated fusion (see Fig 3f).

To assess the leakage free cargo delivery under the assay conditions, we loaded target liposomes with fluorescein (product of the βGlu/FDGlu content mixing assay) and allowed them to undergo fusion with empty ATTO-655-DOPE labeled cargo liposomes with complementary LiNA, see Fig. 3g. (See method for LiNA sequences and conditions). Fig. 3h displays a typical time trace, where cargo liposome docking occurred (signal increase in red channel) but encapsulated fluorescein was not leaked, as shown by the stable signal (blue). 99.2% of the target liposomes (N= 239), remained leakage free throughout the experimental time frame (Fig. 3i), as well as during fusion (Fig. 3g) or upon kiss and run event (see Supplementary Fig. 24 for ensemble measurement).

From the liposome size distribution, the associated distributions of the number of anchored LiNA molecules per target liposome were calculated, assuming complete and homogeneous incorporation^11^. On average, 21.3 molecules of each LiNA (A to F) were anchored to each target liposome, following a lognormal distribution as expected from the liposome size distribution (see Supplementary Fig. 21a). The exact number of LiNAs per individual target will be given by a Poisson distribution around the mean integration. For the smallest 207 target liposomes that underwent fusion in this assay a mean of two LiNA molecules (of each A to F) was calculated, and 30% of those are expected to have one LiNA. For 334 target liposomes a mean of 5 LiNAs (distributed from one to 11) was found. Thus, a low copy number of LiNAs is sufficient for establishing fusion, in agreement with the findings of van Lengerich et al., directly observing fusion in presence of a few LiNA molecules^12^. In SNARE mediated membrane fusion it has been reported that only one to two complexes are sufficient for fusion^36,37^.

### Quantification of high-throughput multiplexing

The breadth and depth of SPARCLD for multiplexing is shown in Fig. 4. Each target liposome is an autonomous nanocontainer, i.e., constitutes an independent experiment, and is stochastically fused with cargo liposomes (A’ to F’) resulting in a distinct sequence of cargo deliveries. Neighboring but otherwise identical target nanocontainers can undergo completely different sequences of fusion events as classified via the distinct fluorescent intensity barcode of the cargo liposomes, see Fig. 4a. The full combinatorial space is multidimensional and cumulatively growing with γ^N^, where γ is the number of cargo populations (each with a unique fluorescent barcode and coupled to a unique LiNA sequence) and N is the number of sequential fusion events per target (see Fig. 4b). A stochastic fusion reaction containing two successive cargo deliveries, results in γ^N^ = 6^2^ = 36 possible permutations. Accumulated with the six permutations with only one fusion event (6^1^) provides 42 possible distinct permutations in total. In our setup with six barcodes, incrementing the number of successive fusions to six or seven will result in >46,000 and ∼0.28 million possible distinct permutations respectively, offering the intriguing possibility of employing SPARCLD for high-throughput analysis of sequential reactions in arrays of immobilized liposomes (see supplementary table 1).

**Figure 4:**
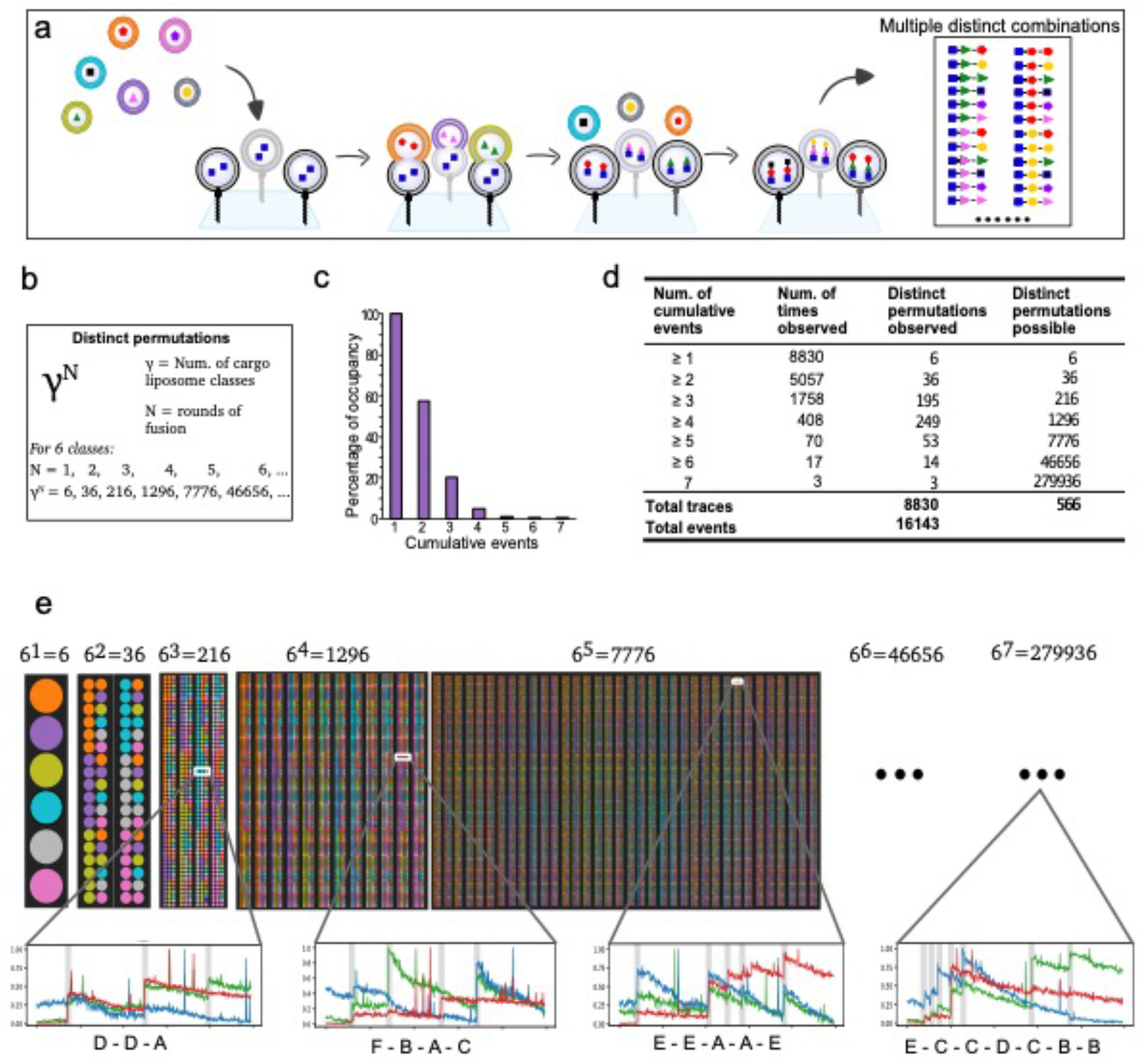
Quantification of High throughput multiplexing. (a) Cartoon representation of possible outcomes of a stochastic combinatorial synthesis of a biopolymer building within each of the atto-to zeptoliter containers. For simplicity a length of three building blocks shown. Each of the parallel tethered target liposomes constitutes an autonomous experiment, where freely diffusing cargos with distinct barcodes and associated LiNA can dock and fuse stochastically. The stochastic leakage free and quantitative fusion can result in combinatorial content delivery effectively turning liposomal nanocontainers into nanoreactors. (b) The method is stochastic both on the order and the number of cargos delivered allowing recording of γ^N^ distinct combinatorial fusions, where γ is the number of LiNA sequences associated with a distinct spectral barcode and N is the number of successive fusions. (c) Bar chart showing the occupancy for one-to-seven rounds of the experimentally recorded successive fusion. (d) Table summing up all observed fusion events, showing successive fusion steps observed on targets, as well as the numbers of times observed, of distinct combinatorial fusions observed along with the possible distinct permutations. The 16,143 total recorded events resulted in a total of 566 distinct sequences. (e) Diagram showing distinct fusion sequences building by multiplexed fusion. The power law increase of possible distinct combinations per fusion round are displayed for up to five rounds displaying 7,776 distinct combinations of cargo delivery. Representative trajectories for both three, four five and up to seven fusion rounds display the distinct sequentially readout and classification from SPARCLD.

To evaluate the operational performance of SPARCLD we recorded more than 8,800 target liposomes, that underwent 16,143 individual fusions events. The high sensitivity of the assay allowed us to directly observe up to seven successive and independent fusion events on individual target liposomes making this, to our knowledge, the most efficient synthetic membrane fusion machinery reported. As expected, the frequency decreased for increasing number of successive fusions (Fig. 4c), probably due to increased electrostatic repulsion by the increasing amount of dsDNA on the surface. Control experiments with non-complementary LiNA, show 4.8±0.9% non-specific binding (NSB) for one fusion, 1.4% NSB for two fusion, and no non-specific interactions above three fusion with 0.9% NSB (see Supplementary Fig. 4). The waiting time distribution of all docking events showed docking to be efficient and occur within the observation time (see Supplementary Fig. 21b). Doubling the amount of LiNAs per liposome resulted in practically identical fusion events (see Supplementary Fig. 25), thus LiNAs were not depleted after the first fusion event, allowing multiple fusions of the same LiNA population. This was further supported by observation of multiple successive fusion events mediated by the same LiNA pair (see Fig. 4e), confirming a non-deterministic fusion, not limited by LiNA depletion.

The possible distinct outcomes and the experimentally observed ones for all one-to-seven fusions rounds are summarized in Fig. 4d. We observed almost all distinct combinations for one-to-three rounds of fusion, confirming the randomization and multiplexing of our method. In 408 cases we found four or more successive fusion events verifying the high efficiency of the LiNA mediated fusion. The large combinatorial space of possible fusion sequences is illustrated in Fig. 4e, albeit limited to five rounds of fusion for clarity, as the possible distinct combinatorial outcome above five rounds becomes too large for informational illustration. Representative time traces for both three, four, five and up to seven rounds of fusion, illustrating the information-rich readout from the real time single particle methodology. The direct real time observation of a total of 566 distinct multiplexed combinations renders SPARCLD a promising method for single particle high throughput multiplexing approaches.

## Discussion

We have developed and validated a platform for the single particle combinatorial lipidic nanocontainer fusion based on DNA mediated fusion (SPARCLD), that allowed the parallelized cargo delivery of sub-attoliter volumes in stochastic order of succession. Nanocontainer annotation relied on a fluorescent barcoding technique based on specific ratios of up to three fluorescent lipids on each liposome. A machine learning framework trained on ground truth barcoded data, accurately and rapidly predicted the barcode identity from three-channel intensity TIRF data and thus the classification of cargo liposomes. Efficient fusion between the immobilized target liposomes was mediated by functionalizing the surface tethered target liposomes with six ssDNA (LiNA) strands and each of the target ones with a complementary ssDNA sequence and a distinct barcode. Real time TIRF-microscopy allows the direct observation of multiple rounds of a highly efficient (93.2 ± 12.0 %) leakage free fusion with content mixing of attoliter volumes for target liposomes on a second timescale and biologically relevant temperature of 37°C.

We have demonstrated spatially resolved and parallel observation of thousands of DNA-mediated single-liposome fusion events. The distribution of different permutations within the characterized combinatorial multistep cargo-to-target fusion sequences revealed that a non-deterministic stochastic delivery of cargo to arrays of immobilized nanocontainers are indeed possible. Each target liposome constitutes an autonomous nanocontainer thus the method allows ∼42,000 target containers per square millimeter (100s of liposomes per field of view) of the microscope surface. SPARCLD quantitative utility is demonstrated for six barcodes combined with six LiNA resulting in parallel recordings of 8,800 individual target containers undergoing more than 16,000 fusion event and resulting in 566 distinct combinations within minutes.

SPARCLD exploits the stochastic sequence of events for high throughput screening and transforms stochasticity from a prohibitive problem in conventional assays into an experimental advantage and an enabling technology for multiplexing. The method can easily be expanded with our 11 complementary LiNA pairs and associated 10 fluorescent RGB barcode libraries available along using our automated ML classification (see SI Supplementary 6 and 7), reaching 10^7^ or up to 10 millions permutations expanding it to High Content Analysis (HCA) providing an ultra-high throughput screening (UHTS) methodology^3^ using picograms of materials^38^. Microfluidic pico-injection integration^39^ or laser microdissection^40^, may expand further sample processing and expand the ultrahigh throughput capabilities.

Being efficient, reproducible and leakage free, we envision SPARCLD can be applied for multiplexed discovery and for combinatorial processing of chemical nanoreactor synthesis for biopolymers such as carbohydrates or nucleic acids reducing material and time cost by several orders of magnitude compared to current state of the art. The combinatorial power of SPARCLD is based on spatially resolved readout and miniaturization. The combination of diameter and thus volume heterogeneities (50-250nm diameters and zepto-to attoliters volumes) with membrane heterogeneities and protein or small molecule partner concentration (10^0^ -10^4^ molecules may create additional distinct combinatorial permutations of regulatory inputs that may be screened with a single molecule readout. We envision this can be combined in DNA-templated synthesis (yoctoliter, single molecule reactors)^41,42^ or used for synthetic biochemical pathways^43^, artificial cells systems^44^ and cell-free expression systems^45^. Integrating SPARCLD with post-combinatorial readout, e.g. for stochastic combinations of protein-ligand interactions, such as for membrane-bound G-protein coupled receptors^46^, and many other drug targets^47^ restricted DNA hybridization and interactions with CRISPR-Cas proteins^18^ could further expand the scope of applications.

## Methods

### Materials

Throughout all experiments milliQ H_2_O (18.2 MΩ, <3 ppm TOC) was used. Reverse-phase and size-exclusion chromatography were carried out on a ThermoFisher Ultimate 3000RS UHPLC system. All experiments are carried out in our buffer HBS500 (HBS, 10 mM 4-(2-hydroxyethyl)-1-piperazineethanesulfonic acid (HEPES), 500 mM NaCl in water, adjusted to pH 7)

### Design, synthesis and characterization of lipidated nucleic acid conjugates (LiNA)

Eleven pairs of 17 *bp* recognition sequences were designed by the following principles to achieve a library with two orthogonal sets: sense-LiNAs (A-L) and antisense-LiNAs (A’-L’), where each pair (AA’, BB’, etc.) has similar binding affinities and lowest possible cross-hybridization. Target liposomes are always engrafted with sense LiNAs, and cargo liposomes always with antisense LiNAs. i) Sense sequences have no C-residues, antisense sequences no G-residues, avoiding any C:G complementarity within each set. ii) All pairs have the same fraction of C:G content, 47.1%, giving a T_m_ above 55 °C, permitting membrane fusion at up to 50 °C. iii) stretches of G-residues were kept to a minimum. iv) Partial complementarity in unwanted combinations kept to a minimum (≤ 10 consecutive bps, and at least 4 nt spaced from the anchor.) v) anchor building blocks for each pair were placed at the 5’- and 3’-end respectively, leading to close proximity between anchors upon hybridization (zipper-like). The anchor-side termini were framed with a non-pairing T-nucleotide giving the strands a charged and bulky structure around the lipophilic part, which disfavors self-aggregation of LiNA strands (e.g., as micelles). Recognition sequences were linked via a triethylene glycol building block which is necessary for high content mixing yields.

**Table S1:**
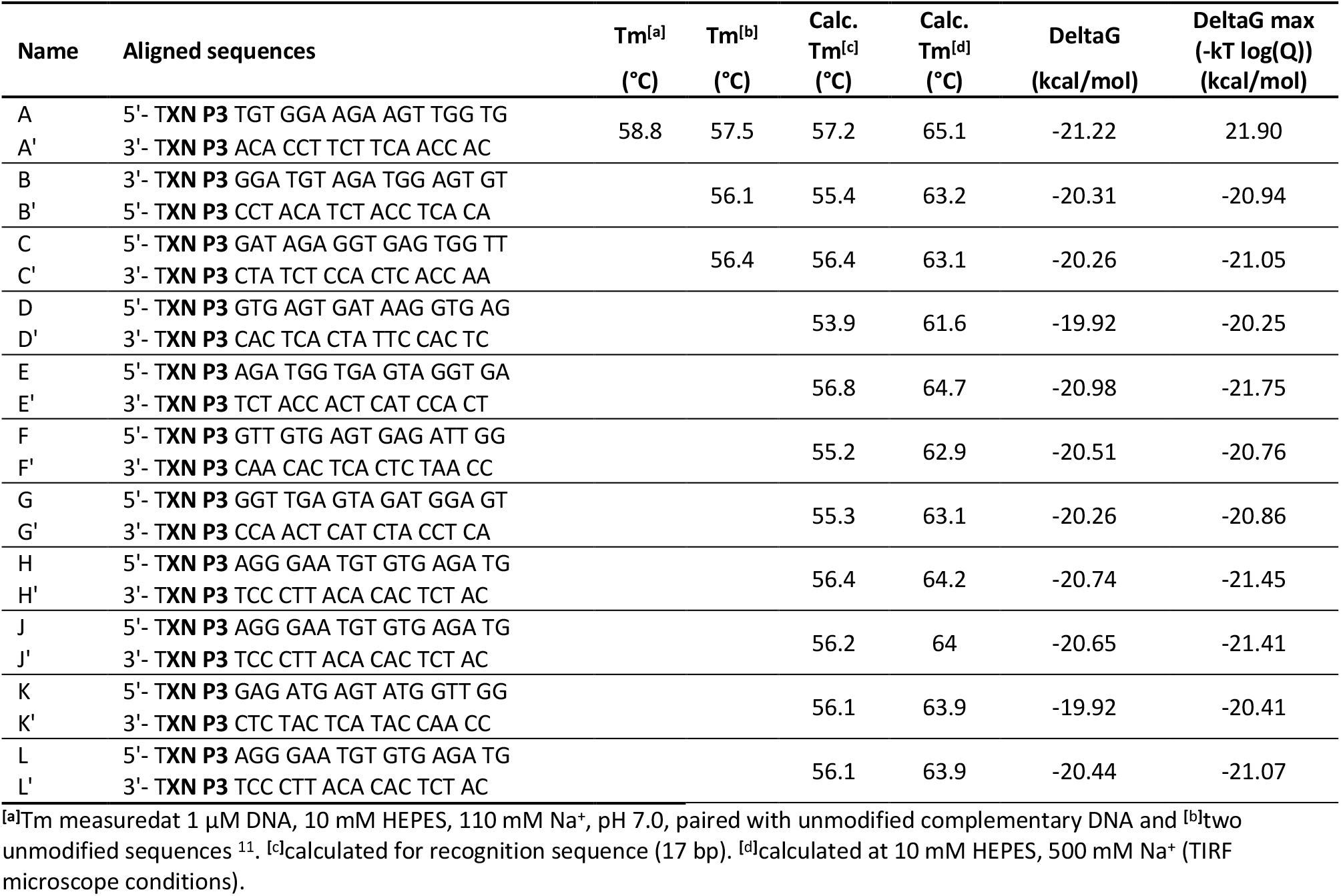
LiNA library with aligned sequences.

The LiNA oligonucleotides were synthesized under standard conditions for solid-phase synthesis. Both the lipid-modification **XN** and spacer **P3** are introduced in the phosphodiester backbone as DMT-protected phosphoramidites. The **P3** spacer is used in a standard solution of 0,1 M in acetonitrile while the **XN** modification is used as a 50 mM solution in a mixture of DCE and acetonitrile (2:1, v:v). Modifications were coupled by hand using a syringe with a mixture of 0.3 mL of the amidite and 0.6 mL of Activator 42, which was flushed through the column twice over a total time of 15 minutes for **P3** and 25 minutes for **XN**. The synthesis and structure of the building block **XN** is previously described ^48^. All oligonucleotides were synthesized without final DMT and cleaved from the solid support with ammonia at 55°C for 16 hours. The solution was filtered, the ammonia was evaporated, and the oligonucleotide was redissolved in 50% acetonitrile/water (v/v). Purification was performed using reverse-phase HPLC (ThermoFisher, Acclaim C8 column, 5 µm, 120 Å), and fractions were tested with MALDI mass spectrometry to verify the purity. The fractions containing product were pooled, tested with analytical HPLC and MALDI, and evaporated to dryness. LiNAs were kept in glass vials at -18°C before the experiments and dissolved in 50% acetonitrile/water (v/v) during the experiments, where it was kept at 4°C. Table S1 summarizes the LiNA sequences used in this study.

**Table S2:**
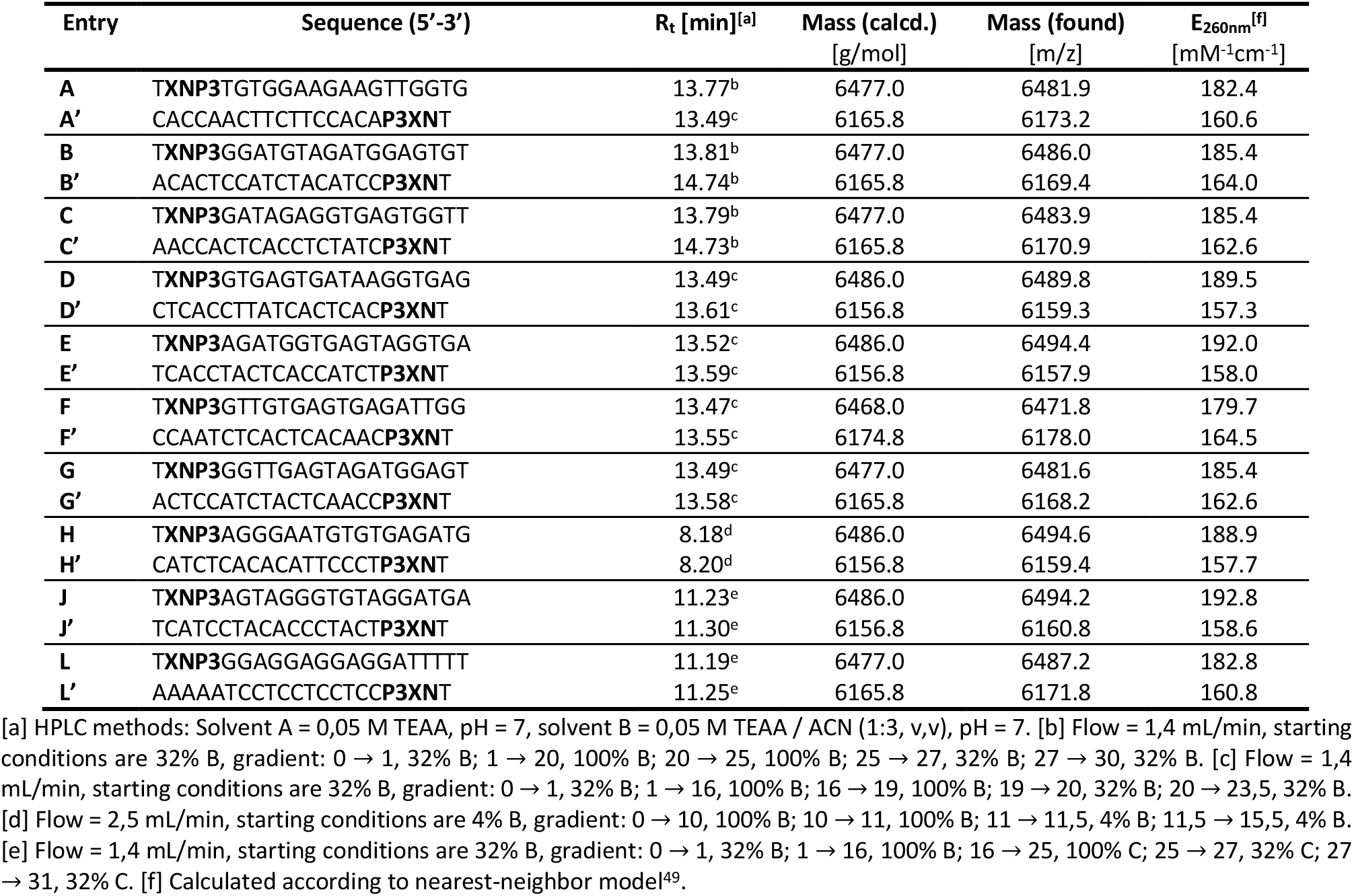
List of LiNA oligonucleotide sequences and their properties.

### Acquisition of TIRF data

All single liposome experiments were accomplished using an inverted total internal reflection fluorescence microscope (TIRF) model IX83 from Olympus. The microscope was equipped with an EMCCD camera model imagEM X2 from Hamamatsu and an 100x oil immersion objective model UAPON 100XOTIRF from Olympus and an emission quad band filter cube, in order to block out laser light in the emission pathway. An incubator system was mounted on the TIRF microscope stage in order to keep a constant temperature at 37°C, and all data was acquired using a 200 nm penetration depth. Three solid state laser lines from Olympus at 488 nm, 532 nm and 640 nm were used to excite DiO, Fluorescein, Alexa 488, ATTO 550 and ATTO 655 fluorophores. For recording of RGB barcoded liposome and multiplexing assay, the emission signals were divided into three channels using a quad band tube turret with dichroic mirrors ZT640rdc, ZT488rdc and ZT532rdc for splitting and with single-band bandpass filters FF02-482/18-25, FF01-532/3-25 and FF01-640/14-25. The image dimension for each channel is 256 times 256 pixels with a dynamic range of 16-bit grayscale. The field of view corresponds to a physical field of view length of 40.96 µm. Recording of content mixing, leakage control and encapsulation efficiency emission signals were guided to the EMCCD camera in bypass mode. The image dimension in these experiments for each channel is 512 times 512 pixels with a dynamic range of 16-bit grayscale. The field of view corresponds to a physical field of view length of 81.92 µm.

### Liposome preparation and LiNA functionalization

SUVs were prepared from mixed lipid films containing 25% cholesterol, 10% DOPE, 1% DOPG charges, 0.1% up to 1.5% fluorescent lipidated labels for the distinct signatures and remaining mole percentage DOPC. The specific barcode lipid mole percentage is as following (all stated in units of mole percentage):

| RGB barcode | ATTO-655<br>DOPE | ATTO-550<br>DOPE | DIO | LiNA functionalization<br>used for multiplexing |
| --- | --- | --- | --- | --- |
| (5,0,0) | 0.5% | 0.0% | 0.0% | A' |
| (0,5,0) | 0.0% | 0.5% | 0.0% | B' |
| (0,0,5) | 0.0% | 0.0% | 0.5% | C' |
| (5,5,5) | 0.5% | 0.5% | 0.5% | D' |
| (5,0,1) | 0.5% | 0.0% | 0.1% | E' |
| (1,0,5) | 0.1% | 0.0% | 0.5% | F' |
| (1,5,0) | 0.1% | 0.5% | 0.0% |  |
| (0,5,5) | 0.0% | 0.5% | 0.5% |  |
| (5,0,5) | 0.5% | 0.0% | 0.5% |  |
| (5,5,0) | 0.5% | 0.5% | 0.0% |  |

0.1% biotinylated lipids were added to the mixture of the target liposomes in order to immobilize on a surface.

### TIRF recording of fluorescent barcoded liposome library for ML classification model

All six SUVs populations were prepared as described in the previous section, but with the addition of 0.1% biotinylated lipids for all six liposome populations with the distinct fluorescent barcodes, to immobilize the liposomes on the neutravidin coated surface. The surfaces were prepared using plasma cleaned Glass slides with fastened sticky-Slide VI 0.4 from Ibidi were functionalized using PLL-g-PEG and PLL-g-PEG-biotin in a 100 to 1 ratio followed by a neutravidin layer as we have done in the past ^18,30^.

Each of the six liposome populations with distinct barcodes were incubated in separate champers on the surface slides for immobilization to achieve vesicle densities of ≈100 vesicles per field of view, incubating for 2 minutes. Inflow of buffer removed unbound freely diffusing liposomes. To compile the specific RGB signature library for model training, liposomes with the desired label concentrations were generated and tethered on passivated surfaces and imaged in identical conditions. (see Supplementary Fig. 11 and methods of liposome preparation and imaging). We tested and trained the algorithm using 10 distinct RGB populations (53,800 liposomes) and selected the best combination offering optimal classification by backward elimination (see Supplementary Fig. 6-9 and Supplementary note 1). The best 6 populations were selected, and more than 100 images of each populations were obtained, corresponding to more than 44.000 where each individual liposome was fitted and colocalized in all three microscope channels and the median of the local background were subtracted. The training library was trained on immobilized liposomes and recorded with the same fast temporal resolution as the dynamic docking and fusion data we want to predict, with all laser lines open at all time (20ms exposure time). Extracted data was used for a supervised machine learning classification, using an extreme gradient boosted decision tree, fitted using the normalized and centered signal ratio in the three channels for all data points.

### TIRF multiplexing assay

Both cargo and target SUVs were prepared as described above and earlier reported^30^. For target SUVs, the suspension contained 0.1% biotinylated lipids for immobilization. Glass slides were prepared and functionalized as described.

Target SUV’s (500 µL, 0.7mg/L) were flushed into the microscope chamber, using a peristaltic pump and left for 5 minutes to immobilize, as to achieve a target density of approximately 70 vesicles per field of view. Remaining freely diffusing targets were washed away by addition of 1 mL buffer, corresponding to approximately five times the chamber volume. The six cargo liposome populations were mixed in an Eppendorf tube with 1.1mL buffer and 0.54mg/L cargo SUV’s of each population and added via a peristaltic pump. The automated TIRF image recording was started, recording 6 fields of view sequentially cycled using automated cellSens imaging software by Olympus. After 4 minutes of recording corresponding to 240 frames, 40 for each field, 0.5 mL of the solution were flown in with a flowrate of 0.5mL/min. The measurement there recorded for a total of 51.14 minutes with 500 cycles, 6150ms/cycle.

### Protein purification

Beta Glucosidase from *Aspergillus Niger* (βGlu) was purchased from Megazymes as a suspension of 3.64 µM enzyme (∼0.44 mg/ml (40 U/ml, 40° C, pH 4.0 on *p*-nitrophenyl β-glucoside, 90 U/mg) in 3.2 M (NH_4_)_2_SO_4_ and stabilized with 0.5 mg/ml BSA. The enzyme was purified as follows. The stock solution was desalted and concentrated using Amicon® centrifugal filter units (MWCO 30 kDa, 3 min per run, 10000 g) as follows: 3 × wash with 500 µl H_2_O, load 500 µl βGlu stock solution, 3 × wash with H_2_O, filling to 500 µl). Retained liquid in filter eluted into a fresh microtube (2 min, 750 g). Two batches of 2 × 500 µl were purified in this manner and pooled to a final volume of 140 µl (∼26 µM βGlu) and 125 µl (∼29 µM βGlu, respectively, and transferred to HPLC insets. (∼3.52 mg/ml). The concentrated enzyme mixtures were then purified using size-exclusion chromatography (Agilent AdvanceBio SEC, 2.7 µm, 300 Å, 150×7.8 mm, fractionation range 5 – 1250 kDa). Isocratic method: 15 mM phosphate buffer, 140 mM Na^+^, pH 7.4 (P/Na), 0.35 ml/min, 19 min, 1.7 nmol per injection. βGlu was collected based on UV absorbance at 260 nm (Ret. time 8.8 ± 0.2 min) in four fractions (200-250 µl). Activity for each fraction (25-fold diluted) was measured against a standard curve of diluted stock solution of the enzyme (0.5 – 4 U/ml) in HBS based on a fluorimetric assay on the conversion of fluorescein di-β-D-glucopyranoside (FDGlu, 0.1 µM) to fluorescein (Varian Cary Eclipse, Excitation/emission 488/510 nm; temperature controller, 37 °C; PMT, 800V; excitation and emission slits, 5 nm; 250µl sample per cuvette). The enzyme concentration correlated linearly with the intensity after 20 min reaction time at 37 °C. All HPLC fractions were pooled and the volume adjusted to 1 ml with a final concentration of 6.4 µM (6.4 nmol, 0.77 mg/ml) βGlu.

### Fluorescent labeling of protein

Purified β-Glucosidase (3.2 nmol) was fluorescently labelled with a 30-fold excess of Alexa Fluor™ 488 sulfodichlorophenol ester (10 mM stock solution in anhydrous DMSO) to ensure labeling of all enzymes. To treat both labeled and unlabeled enzymes equally, two aliquots of 500 µl of enzyme solution were treated in parallel. First, both aliquots were concentrated approx. 10-fold by a single pass through a centrifugal filter (as above, 5 min, 10000 g) to 50-52 µl each and eluted into a fresh vial. The pH in both vials was adjusted to ∼8.3 by addition of 2.5 µl 0.2 M Na NaHCO_3_ (pH 9.0). Then, the appropriate aliquot of dye was added to *one* of them. After 1 hour of incubation at room temperature, both samples were purified using a size-exclusion spin column (Illustra® microspin S-200HR, GE-healthcare) to exchange the buffer and remove unreacted dye. After pre-equilibration with HBS500 (3 × 200 µl, passed through the column bed by centrifuging at 700 g for 1 min). The samples were loaded (∼55 µl) and eluted into a fresh vial (700 g, 2 min) and the column was washed with 50 µl HBS500 and collected into the same vial (2 min, 700 g). The eluted enzyme was now suspended in 100 µl HBS500 at a final enzyme concentration of min. 26 µM (32 µM based on volume; multiplied by the recoveries stated by the supplier – spin filter: ≥95%, size-exclusion ≥85%).

### TIRF substrate leakage control

Target liposomes were prepared as described with addition of 0.1% biotinylated lipids and no membrane fluorescent markers were added and functionalized using 1:500 LiNA to lipid ratios. 500 µM fluorescein was encapsulated during rehydration following recently published methodology ^28,51^. Cargo liposomes were prepared with addition of 0.5% ATTO 655 DOPE and functionalized using 1:2000 LiNA to lipid ratios, complimentary to the targets using LiNA D for targets and D’ for cargo vesicles. Both populations were exposed to 10 cycles of flash-freezing and thawing as to ensure an unilamellar membrane structure followed by extrusion at 100nm. Target SUV’s were flushed into the assay using a peristaltic pump and allowed to immobilize on the surface, as to achieve a target density of approximately 300 vesicles per field of view. Excess target liposomes were washed thoroughly away along with excess freely non encapsulated fluorescein. Images were acquired for 400ms using 488nm laser and 100ms using 640 nm laser, using the two laser lines sequentially. 4 positions were recorded in 40 cycles and a 10 second change time, providing a temporal resolution of 42.079 sec between per cycle. Image analysis quantification of docking and leaking was done using homemade software in python.

### TIRF encapsulation efficiency control

Target liposomes were prepared as described with addition of 0.1% biotinylated lipids and 0.1% ATTO 655 for membrane fluorescent marking and rehydrated in the presence of 3.2 nmol Alexa Fluor™ 488-labeled ß-Glucosidase (26 µM protein, see above). The proteins were encapsulated during rehydration with a final concentration of 32 µM following recently published methodology ^28,51^. The flash-freezing cycles were reduced to 3 as to minimize denaturation of the proteins. Liposomes were extruded at 100 nm.

Evaluation and correction of potential intensity crosstalk in the microscope channels was done by assembly of otherwise identical target liposomes were prepared without encapsulation of the protein for membrane signal crosstalk subtraction. Target SUV’s were flushed into the assay to immobilize with a density of approximately 300 vesicles per field of view. 100 images were recorded of both the protein encapsulated liposome population and the empty control liposome population.

### TIRF content mixing quantification assay

Target liposomes were prepared as described with addition of 0.1% biotinylated lipids and 0.1% ATTO-550-DOPE for membrane fluorescent marking. 3.2 nmol purified ß-Glucosidase were encapsulated during rehydration with a final concentration of 32 µM as described in the previous section. The flash-freezing cycles were reduced to 3 as to minimize denaturation of the proteins followed by extrusion at 100 nm. Cargo liposomes were prepared as described and were membrane labeled with 0.1% ATTO-655-DOPE. 250 µM Fluorescein di-β-D-glucopyranoside (FDGlu) was encapsulated during rehydration and exposed to 10 cycles of flash-freezing and thawing followed by extrusion at 100 nm. The liposomes were used the same day as preparation, LiNA-functionalized shortly after preparation and only diluted immediately prior to their measurement (889 - fold dilution, resulting in a final cargo liposome concentration of 0.0045g/L). Target liposomes were functionalized using 500:1 lipid to LiNA sequence D. Functionalization of cargo vesicles were done using 2000:1 lipid to LiNA sequence D’ for complimentary specific interaction and D for non-complementary controls.

Target SUV’s were flushed into the assay to achieve a target density of approximately 300 vesicles per field of view, and excess target liposomes were washed thoroughly away. Images were acquired for 100ms for each of the three channels, using all three laser lines sequentially, with 4 field of view positions. 300 cycles of the 4 positions were recorded with a temporal resolution of 21.352 sec per cycle. Cargo liposomes were flushed into the image chambers after 21 frames corresponding to 7.2 minutes with a flow of 0.5ml/min.

### Liposome preparation for bulk assays

Target and cargo liposomes were prepared with the same lipid composition as for the TIRF content mixing quantification assay, based on 2 µmol total lipid. Rehydration of lipid films: *target*, labelled with 0.1% biotinylated lipids and 0.5 mol% ATTO-550-DOPE were rehydrated 26 µM βGlu in HBS (30 min RT, then 3 × freeze/thaw); *cargo* and *control*, labelled with 0.5% ATTO-655-DOPE, in 250 µM FDGlu substrate or pure HBS, respectively (both 30 min 50 °C, then 10 × freeze/thaw). Final lipid concentration: 20 mM. The suspensions were extruded through 100 nm pores (double-membrane, 10 passes in the same direction, customized hand extruder). Unentrapped βGlu and FDGlu were removed from the target and cargo liposomes, respectively, using size-exclusion HPLC, as described, but carried out at 20 °C and using fluorescence detection to identify liposome-containing fraction. The resulting liposome concentration was based on the dilution during chromatography, typically 0.8-1.1 mM.

### FDGlu leakage control

Immediately after purification, cargo liposomes were engrafted with LiNA B’ at 0.2 mol% (Lipid/LiNA ratio 500:1) and incubated for 30 min at room temperature. Leakage of entrapped FDGlu was then measured in triplicate, based on the fluorescein production by externally added β-glucosidase as measured by time-course fluorescence spectroscopy. In brief, FDGlu-entrapping liposome suspensions (intact 100 µM total lipid, in 250 µl HBS) were either lysed (25 µl, 1% w/w Triton-X 100) or left intact (25 µl buffer). Then β-Glucosidase (22 µg/ml, 0.18 µM) was added while monitoring the fluorescence (Ex. 488, Em. 510, same settings as for activity assay, above) for up to 24 h at 37°C. As the enzyme cannot pass the lipid bilayer, it can only convert leaked/lyzed substrate. The percentage of leakage (%L) was determined as: %L = (I_intact_-I_0,intact_)/ (I_lysed_-I_0,lysed_).100%. After 24h still less than 50% of FDGlu had leaked.

### Fusion assay in bulk

Immediately after purification, the liposomes were engrafted with LiNAs and all incubated at room temperature for at least 15 min. Fusion experiments were carried out with i) complementary LiNAs with substrate (*target*: LiNA-D, *cargo*: LiNA-D’) ii) non-complementary LiNAs with substrate (LiNA-D on both) or iii) complementary LiNAs but without substrate (*target*: LiNA-D, *control*: LiNA-D’ (Lipid:LiNA ratio 500:1, all incubated at room temperature ≥15 min). Measurement conditions: The Förster Resonance Energy Transfer (FRET) channel (Ex. 532 nm, Em. 680 nm) between the ATTO-550-DOPE (*target*, donor) and the ATTO-655-DOPE (*cargo*, acceptor) was monitored. Fusion between *target* and *cargo* liposomes mixes the lipids between the differently labelled membrane, increasing the FRET between the labeled lipids. After establishing a baseline signal with *target* liposomes a 1:1 aliquot of 100 µM }bGlu} (final: 50 µM lipid in each population) added and the next data-point recorded as quickly as possible (t_0_, within 3s after mixing) with the temperature controller set to 37 °C (preheated cuvettes), running in parallel conditions i) through iii) in a total of four replicates (Supplementary Fig. 22). At the end of the experiment, 0.1% Triton X-100 was added to ensure that lysis would disrupt the observed FRET, ruling out direct interactions between the FRET dyes

### Tracking and co-localization software for TIRF multiplexing experiments

In order to track and localize target liposomes and co-localize the position in all imaging channels we used in-house developed python software, as to ensure a nanometer precise colocalization in all three colors throughout the entire measurement, even with the usage of a continuous flow introduces by the peristaltic pump. The tracking and localization software is used and published in earlier publications ^30,52^. The developed methodology localizes all target SUVs on a surface using TrackPy and subsequently collects the signal from each spot and corrects for background noise throughout the experiment, returning a time trace of intensities for all three channels for each target.

### Liposome docking and signal convolution

Time trajectories after tracking and colocalization were normalized and analyzed using a digital signal convolution algorithm developed for this study. The algorithm convolves each normalized trajectory individually for all three channels with an idealized fusion step function for identification of fusion. The product of the single-step signal from fusion of a cargo liposome together with a step-function results in peaks at the exact arrival time, as seen in the lower trace from Fig. 2c (see Supplementary Fig. 10). A third power function applied on top of the convoluted signal enhances it, eliminating false docking detection, and peaks detected using the SciPy signal processing package. The raw intensities for identified fusion were found and classified using the supervised machine learning model returning the predicted barcode for each identified fusion step.

## Acknowledgements

We thank K.J. Jensen for useful discussions.

This work was funded by the Villum foundation by being part of BioNEC (grant 18333) for M.G.M, P.M.G.L, N.A.R., S.V., and N.S.H. Villum foundation young investigator fellowship (grant 10099), and the Carlsberg foundation Distinguished Associate professor program (CF16-0797), and the NovoNordisk Center for Biopharmaceuticals and Biobarriers in Drug Delivery (NNF16OC0021948) for N.S.H. Work at The Novo Nordisk Foundation Center for Protein Research (CPR) is funded by a generous donation from the Novo Nordisk Foundation (Grant number NNF14CC0001).

N.S.H. is a member of the Integrative Structural Biology Cluster (ISBUC) at the University of Copenhagen.

## Author Contributions

M.G.M, S.S-R.B, P.M.G.L and N.S.H wrote the paper with feedback from all authors. S.V and P.M.G.L designed the LiNAs and N.A.R synthesized the sequences and P.M.G.L performed all the characterization measurements in bulk. M.G.M designed, carried out and analyzed all TIRF microscopy experiments, and prepared all liposomes, trained the Machine learning algorithm. M.B.S implemented the Machine learning algorithm together with M.G.M. S.S-R.B and M.G.M wrote the automated tracking and event finding analysis. M.G.M and P.M.G.L with inputs from S.V and N.S.H, planned the encapsulation and content mixing liposome assays which were carried out by M.G.M and P.M.G.L. S.B.J and M.Z helped with imaging and analysis. P.H. helped in analyzing the data. N.S.H conceived the project idea in collaboration with S.V. and had the overall project management and strategy.

## Corresponding Authors

Correspondence should be addressed to Nikos S. Hatzakis., or Stefan Vogel.

## Supporting information

Is available for this paper.

