## Supplementary materials for "Single particle combinatorial multiplexed liposome fusion mediated by DNA"

### Table of content

**SI Fig. 1** – Zipper-like membrane fusion facilitated by complimentary LiNA stands 2
**SI Fig. 2** – Calculation of LiNA hybridization in multistrand mixtures 3
**SI Fig. 3** – Suppression of non-specific binding to the passivated surface 4
**SI Fig. 4** – Quantification of the total number of specific subsequent docking and fusion events as well as the respective ad control for non-complimentary LiNA mediated fusion 5
**SI Fig. 5** – Representative traces and the associated signal convolution and ML predictions for accurate barcoding classification 6
**SI Fig. 6** – Intensity in all 3 fluorescent channels for the 10 populations making up the barcoding library 7
**SI Fig. 7** – Prediction accuracy for the 10 populations making up the barcoding library 8
**SI Fig. 8** – Trees from forward and backward selection experiments 9
**SI Fig. 9** – Confusion matrices displaying classifications accuracy for varying number of barcode populations. 10
**SI note 1** – Selection of optimum subset library of barcoded liposomes 11
**SI Fig. 10** –Signal convolution algorithm with idealized step for docking event finding 12
**SI Fig. 11** – Reproducibility in preparation of liposomes, 3 biological repetitions and imaging by two independent persons 13
14
**SI Fig. 12** – Ground truth data validation of the machine learning model performance with liposomes associated with LiNA D' and fluorescently barcoded (5,5,5). 14
**SI Fig. 13** – Purification of β-glucosidase 15
**SI Fig. 14** – Example size exclusion of liposomes encapsulating βGlu 16
**SI Fig. 15** – Example size exclusion of liposomes encapsulating FDGlu 17
**SI Fig. 16** – Liposome characterization using Nanoparticle Tracking Analysis (NTA) 18
**SI Fig. 17** – Representative Trajectories of successful cargo delivery by DNA mediated fusion 19
**SI Fig. 18**– Representative Trajectories of control experiments displaying liposome fusion for empty cargo liposomes with complimentary LiNA sequences showing no product formation 20
**SI Fig. 19** – Representative Trajectories of control experiments of cargo liposomes with non-complementary LiNA sequences displaying no prolonged docking and thus no fusion events 21
**SI Fig. 20** – Relation between cargo liposome size and target liposome size 22
**SI Fig. 21**– Quantification of Docking lag time and fusion for individual LiNA sequences 23
**SI Fig. 22** - Lipid mixing monitored by FRET in spectrometer 24
**SI Fig. 23** – Encapsulation efficiency of β-glucosidase 25
**SI Fig. 24** – Leakage of FDGlu substrate recorded by spectrometer 26
**SI Fig. 25** – Number of docking and fusion events are not limited by LiNA depletion 27
**SI table 1** – Possible distinct permutations for different sizes of cargo libraries for multiplexing assay 28

**REFERENCES** **28**

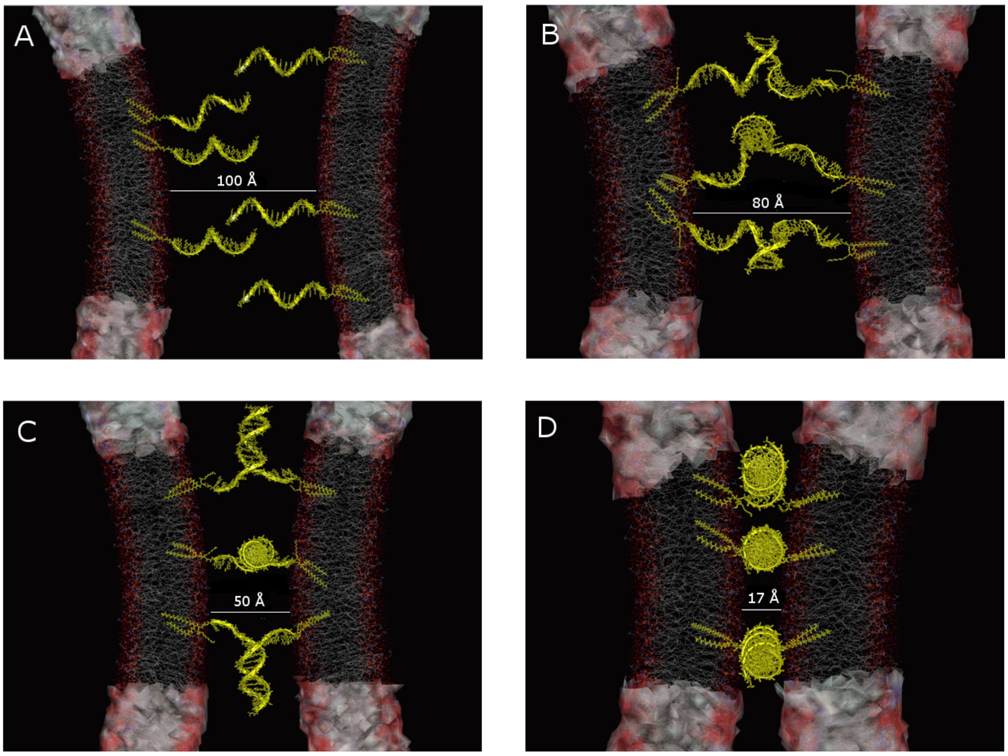


#### Supplementary Figure 1 – Zipper-like membrane fusion facilitated by complimentary LiNA stands

Complementary LiNA sequences are designed to operate like a zipper with the LiNA on the target and thus facilitate a highly efficient zipper-like fusion of membrane. The zipper mechanism will, upon docking of cargos functionalized with LiNA, zip with the complimentary LiNA on the target liposome and facilitate close membrane contact enabling the lipid mixing followed by fusion and thereby content mixing. Figure A to D shows simulated distances for the zipping mechanism[1]. The speed of fusion is found to increase with temperature, with activation energies for fusion with vesicles based on DOPC on 30kBT (34.3±0.8 kBT)[2]. For comparison our system is based on 60 mole % DOPC lipids. Our system relies instead upon a lower biological relevant temperature, 37 degrees Celsius, as to enable the study and multiplexed screening of biological assays. The LiNA sequences two $C_{16}$ hydrocarbon chains spontaneously incorporate into the lipid bilayers as we showed recently[3].

| **Equilibrium binding** | | LiNA on incoming cargo liposome | | | | | | |
|---|---|---|---|---|---|---|---|---|
| | | A' | B' | C' | D' | E' | F' | H' |
| Bound to | A | 0.9998 | 0.0000 | 0.0000 | 0.0000 | 0.0000 | 0.0000 | 0.0019 |
| | B | 0.0000 | 0.9996 | 0.0000 | 0.0000 | 0.0000 | 0.0000 | 0.0051 |
| | C | 0.0000 | 0.0000 | 0.9992 | 0.0001 | 0.0040 | 0.0001 | 0.0085 |
| | D | 0.0000 | 0.0000 | 0.0000 | 0.9981 | 0.0012 | 0.0002 | 0.0080 |
| | E | 0.0000 | 0.0000 | 0.0007 | 0.0010 | 0.9940 | 0.0000 | 0.0011 |
| | F | 0.0000 | 0.0000 | 0.0000 | 0.0005 | 0.0100 | 0.9995 | 0.1850 |
| | unbound | 0.0002 | 0.0004 | 0.0001 | 0.0003 | 0.0005 | 0.0002 | 0.7801 |

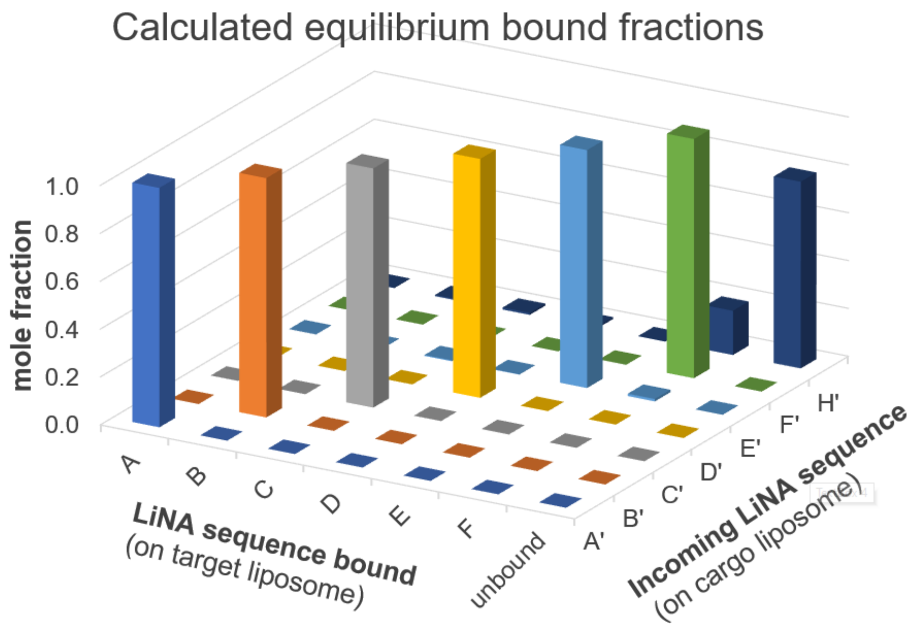


#### Supplementary Figure 2 – Calculation of LiNA hybridization in multistrand mixtures

Theoretical analysis representative of the binding of the LiNAs of an incoming cargo liposome (A’-F’, control = H’) to the available LiNAs on the target liposome (A-F) or remaining unbound. Calculated using the NUPACK web application[8], conditions: All DNA strand conc. 1 µM, 0.51 M $Na^{+}$, 37 °C, matching TIRF assay conditions. The table shows the individual LiNA pair mole fractions of at least $10^{-4}$. The 3D graph illustrates the same numbers. The control sequence H’ was included, as it was used as a non-complementary control in the combinatorial TIRF setup. It is exhibiting low non-specific binding to F’, which could explain some of the docking events in the control experiment.

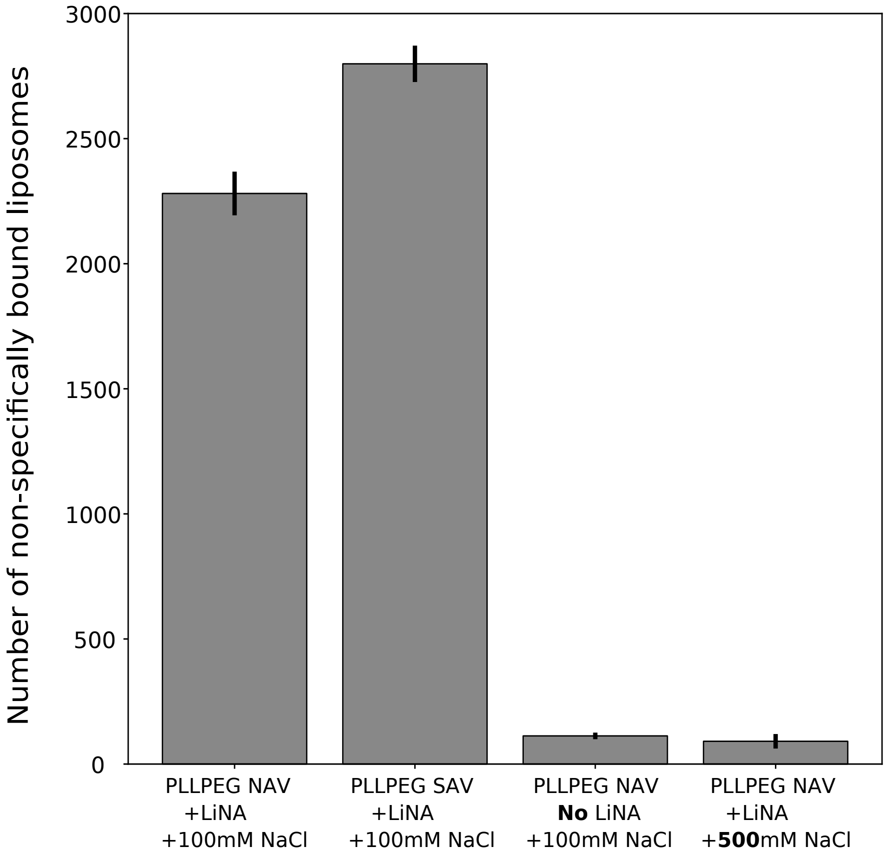


#### Supplementary Figure 3 – Suppression of non-specific binding to the passivated surface

Surface passivation techniques effect to minimize non-specific binding of cargo liposomes on the surface. LiNA engrafted liposomes display high degree of non-specific binding for PLLPEG passivated surfaces coupled with either Neutravidin or Streptavidin. Non-specific binding is minimal for liposomes not loaded with LiNA hence non-specific binding to PLL-PEG passivated surface is driven by LiNA on liposomes, possibly via charges. Increasing NaCl concentration from 100mM to 500mM practically eliminated non-specific binding of LiNA loaded liposomes to passivated surface. The addition of 500mM NaCl minimized non-specific binding to non-specific binding found for non-DNA coupled liposomes were tested combined with both Neutravidin to facilitate the irreversible immobilization of target liposomes on the surface. From the analysis we find very low nonspecific interactions of liposomes with PLL-PEG passivation, but LiNA functionalization on the liposomes increases the non-specific interactions possibly due to charges. This can be suppressed by using Neutravidin for immobilization and passivation on the surface, along with increased ionic strength (NaCl).

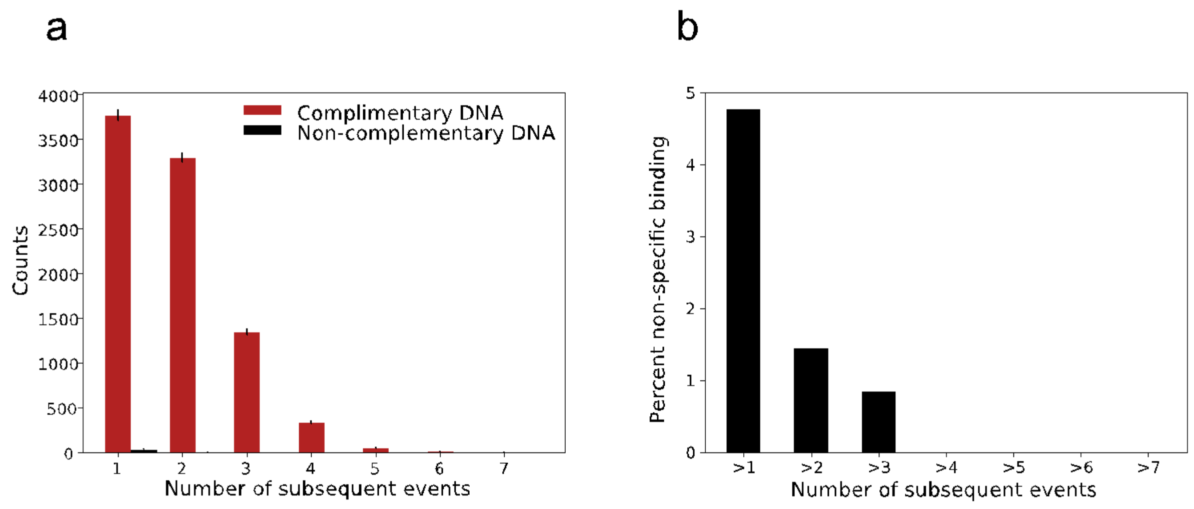


Supplementary Figure 4 – Quantification of the total number of specific subsequent docking and fusion events as well as the respective ad control for non-complimentary LiNA mediated fusion

a) Red Bars: Number of fusion events divided into the number of subsequent fusion events per target liposomes. Data for specific interactions driven by both complementary LiNA. Black bars correspond to non-specific for liposomes loaded with non-complementary DNA. b) We analyzed the low number of non-specific events from Supplementary Fig. 4a, by normalizing the data with the number of experiments. The non-specific binding of one or more subsequent fusion per target liposome constitutes 4.8±0.9% of the total events. Analyzing target liposomes undergoing two or more subsequent fusion events, the non-specific binding decreases to 1.4%, for targets undergoing three or more we see 0.9% non-specific binding which decreases to 0% analyzing four and above subsequent events.

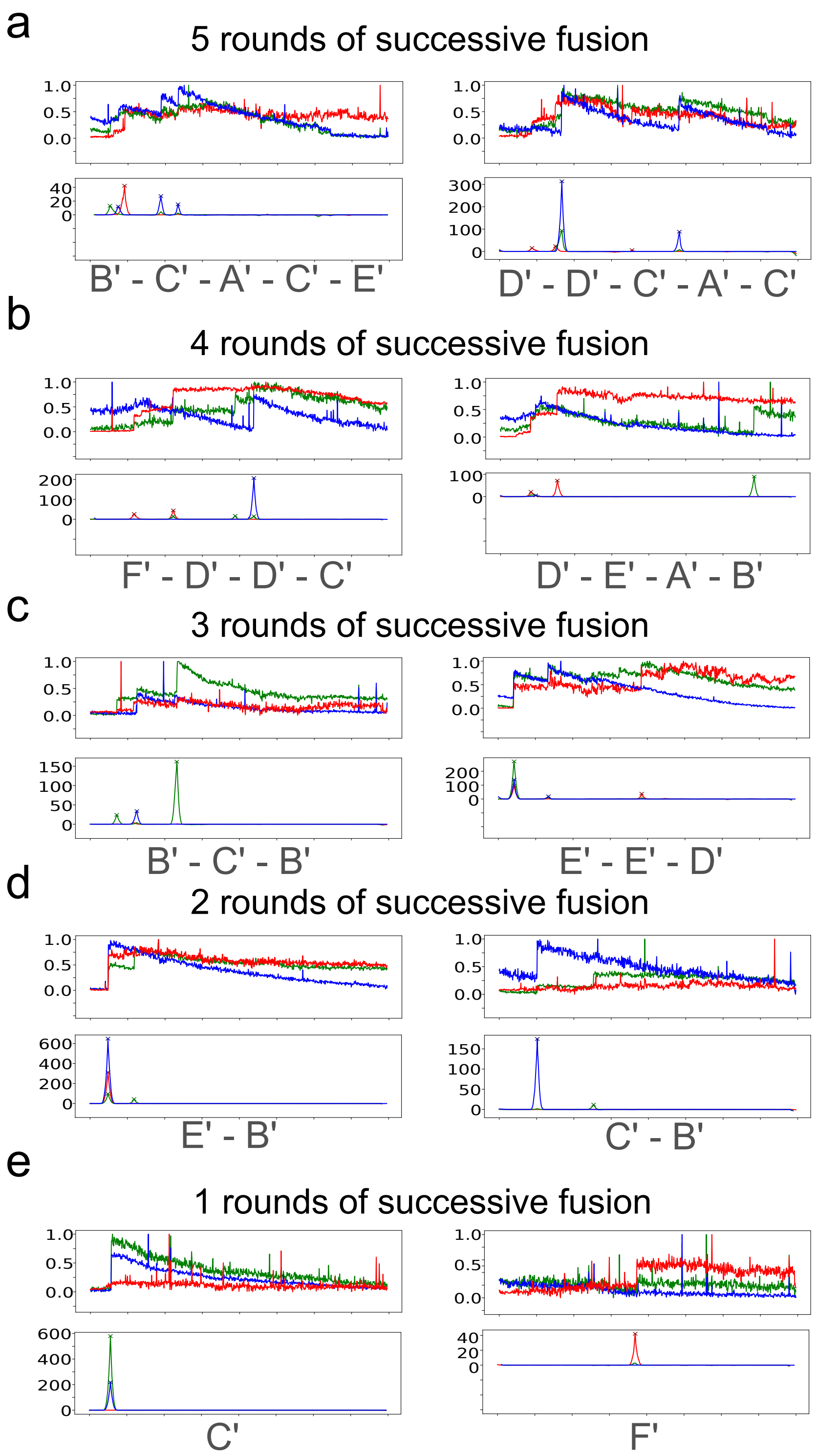


#### Supplementary Figure 5 – Representative traces and the associated signal convolution and ML predictions for accurate barcoding classification

Representative time trajectories displaying 1 to 5 successive docking events. Note tracked target liposomes can either transiently interact with cargo liposomes kiss&run events or irreversibly dock for prolonged time leading to fusion. Nanocontainer annotation relies on a fluorescent barcoding technique and Signal convolution for docking. Letters correspond to the barcoded liposomes identity and distinct LiNA sequences thereof.

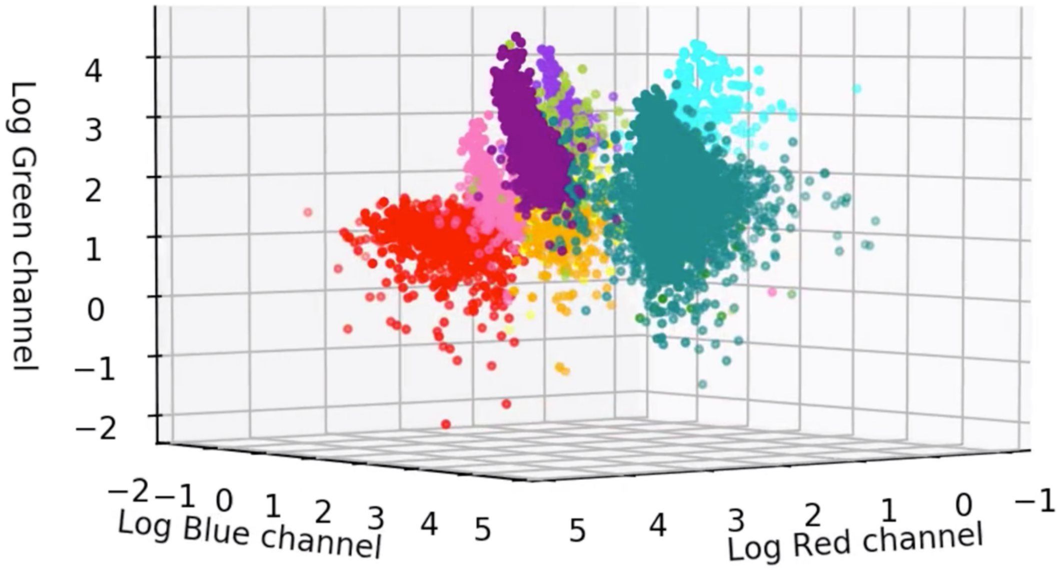


Supplementary Figure 6 – Intensity in all 3 fluorescent channels for the 10 populations making up the barcoding library

Liposomes with multiple distinct combinations of three fluorescently labelled lipids were designed, tested and classified. 10 of the combinations demonstrated distinct combinations of emission in the three microscope channels. Each individual liposome was colocalized and background subtracted in all three channels. A supervised machine learning model was trained on the ratios for recognition and classification Here we show the logarithm of the intensity ratio in all three channels.

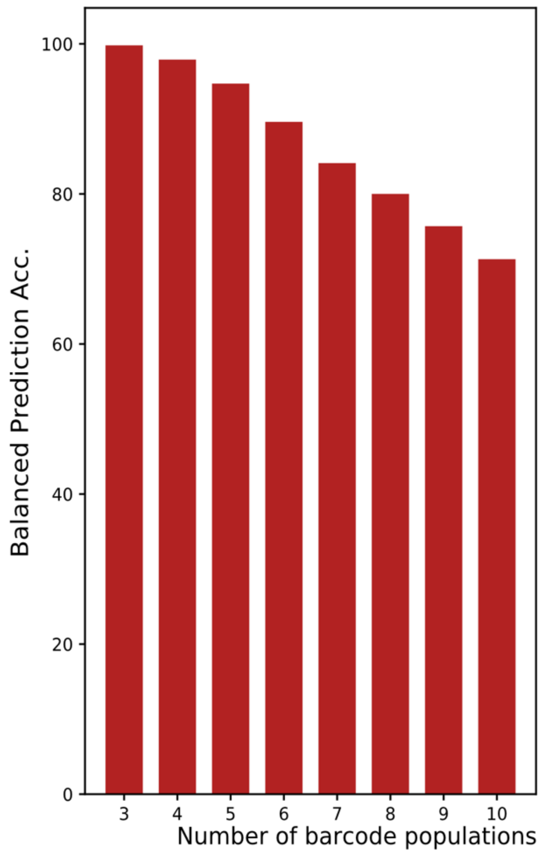


Supplementary Figure 7 – Prediction accuracy for the 10 populations making up the barcoding library

Balanced prediction accuracy for three and up to 10 liposome classes. The balanced accuracy score is defined as the average of recall obtained on each class. The error/confusions are stated from individual the confusion matrices in supplementary figure 5 for each class up to 6 classes. The bar plot shows that for a larger dataset with multiple classes for recognition, the machine learning prediction holds.

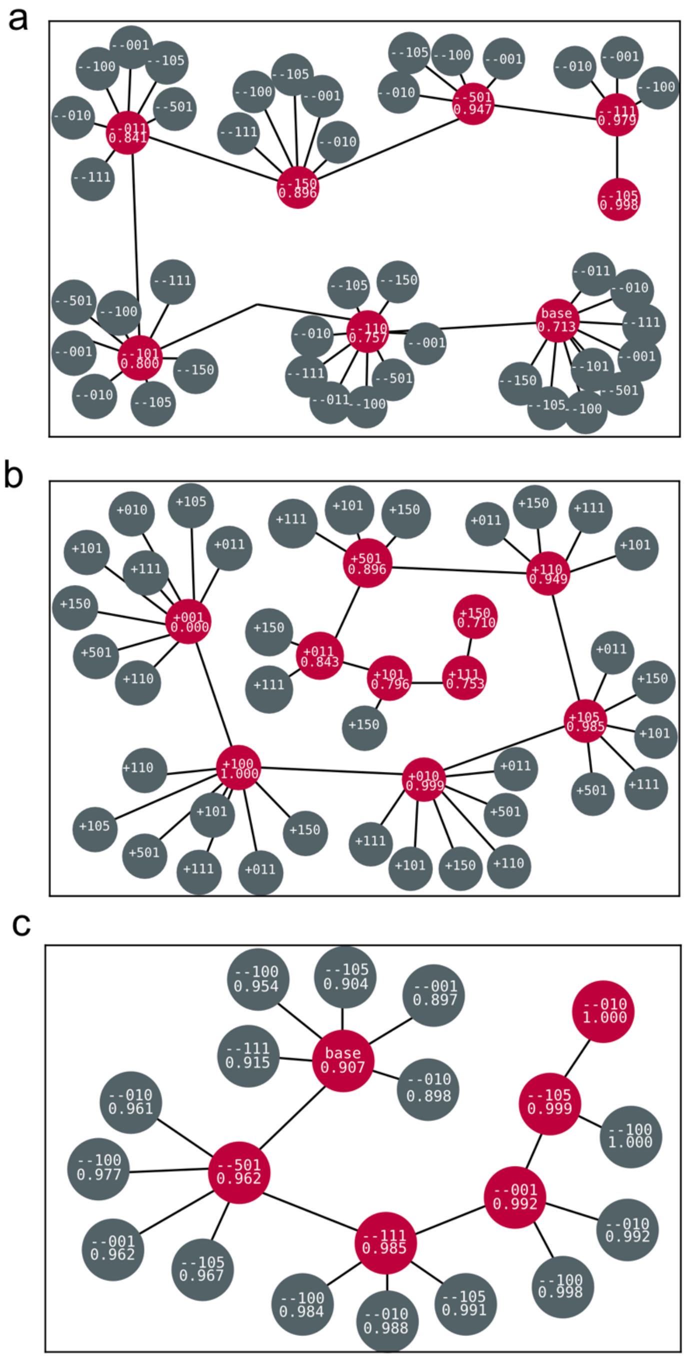


#### Supplementary Figure 8 – Trees from forward and backward selection experiments

a) Backward selection tree, where each node represents a unique subset of the data. The 'base' node contains all 10 populations, and the excluded population for each step can be seen in the label of the corresponding node, along with the corresponding balanced accuracy of the trained model. Red nodes represent the nodes selected during each round of deselection, while grey nodes represent the nodes that were not selected. b) Forward selection tree, where each node represents an additional population of liposomes added to the analysis, along with the corresponding accuracy. The results of a) and b) are summarized in the Supplementary Table 2. c. Backward selection, as in a) for the 6 liposome populations used in the fusion assays. Balanced accuracy scores are provided for all models.

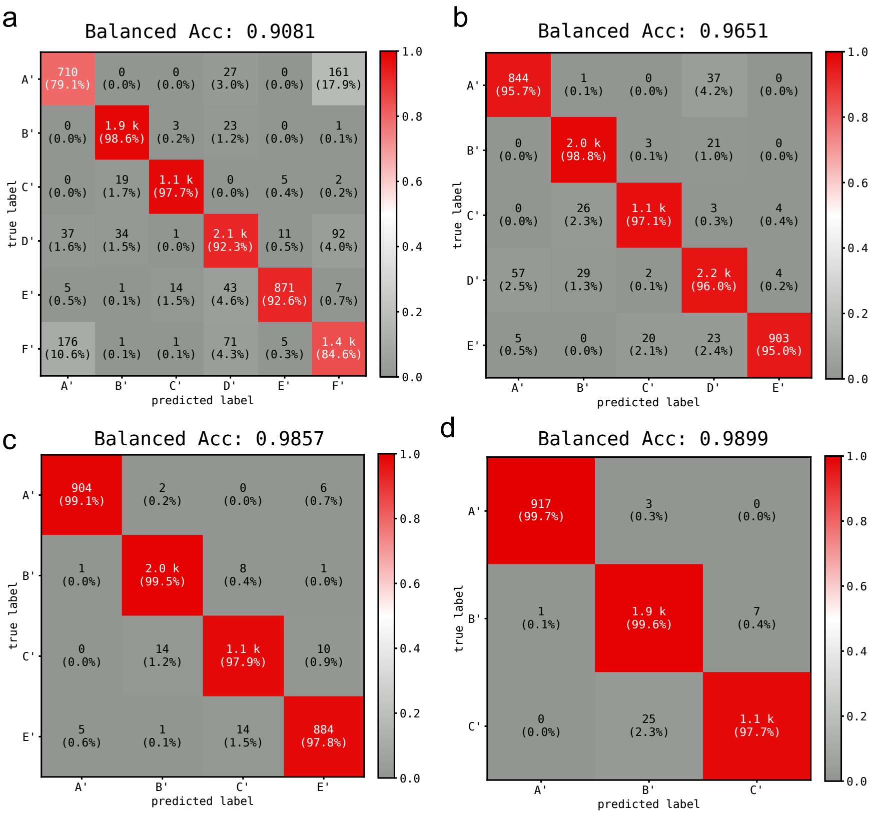


#### Supplementary Figure 9 – Confusion matrices displaying classifications accuracy for varying number of barcode populations.

a) Confusion Matrix for the complete model ('base') in Supplementary Fig.6c. Top number in each cell corresponds to the absolute number of selected data points, the parenthesized number represents the row-wise percentage of selected data points. The diagonals represent the correctly classified data, where the diagonal percentages represent the class-specific recall as a percentage. b)-d) Confusion Matrices for subset models with one, two, and three populations excluded, respectively. Removal of the most difficult to discern population by backward selection results in improved balanced accuracy. This also supports the argument for prioritizing the rank from backward selection experiments, as the recall for the LiNA A' population in subfigure B rises drastically.

#### Supplementary note 1 – Selection of optimum subset library of barcoded liposomes

Tree boosting methods are ubiquitous within Data Science and Machine Learning communities as out-of-the-box, scalable classifiers that have documented high performance across different types, and scales of data. In order to classify different barcodes of liposomes we used XGBoost, a boosted tree system for training boosted tree-based classifiers rapidly on large datasets. We chose to utilize XGBoost as it scales efficiently on large, sparse datasets, and due to the fact that it minimizes softmax-categorical cross entropy during learning, which typically serves as the loss function for categorical data for both tree-based methods and state-of-the-art deep neural networks [4]. We further draw advantage from the fact that boosted trees are linear classifiers, so minimal preprocessing and feature engineering is needed on raw data to achieve state-of-the-art results.

In order to categorize the multiplexing events with maximum accuracy, we produced 10 distinct populations of barcoded liposomes, as described in Supplementary Table 2. We selectively trained XGBoost models with subsets of the populations, using a forward/backward selection scheme, loosely inspired by the methodology described in Hastie. Doing so permits us to choose the best subset of models and allows us to find the combination of populations that result in the most accurate predictions. We base our scorings on the balanced accuracy, which is defined as the average recall across all classes.

**Preprocessing**

Background-corrected, three-channel intensity data of barcoded liposomes was preprocessed by calculating the ratios of intensities between the three channels separately.

**Methodology for forward and backward selection**

For the backward selection experiment, we trained an initial model on the full set of 10 populations. For each round of deselection, a model was trained for each subset with one population excluded. Of the resulting 10 subsets, the one with the highest balanced accuracy score was passed on to the next round of selection. The rounds of deselection continue until the specified number of populations remain.

For the forward selection experiment, we started with the 0:0:5 (single color) population, of liposomes. We train one model for each of the remaining populations and choose the population which resulted in the highest balanced accuracy score. The rounds of selection continue until the specified number of populations are selected.

**XGBoost Hyperparameters**

We modified the standard implementation of XGBoost with the SkLearn API as described in table Supplementary Table 2 {...}[5]. All models with three or more populations were trained to minimize categorical cross entropy, while models with only two populations were trained to minimize binary cross entropy.

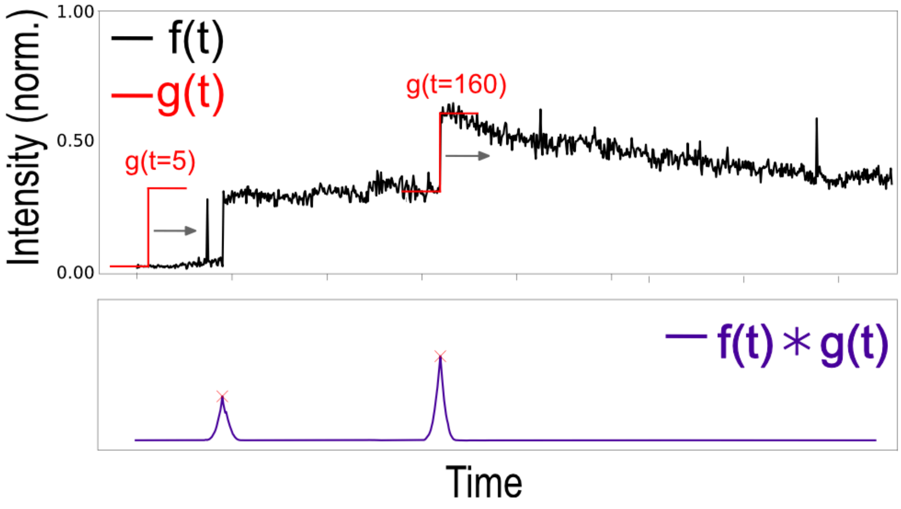


Supplementary Figure 10 –Signal convolution algorithm with idealized step for docking event finding

Each target liposome is tracked and colocalized in the three microscope channels, giving three time series trajectories for each target. These three trajectories are normalized, and individually analyzed. f(t) is a representative trajectory, showing some spikes (kiss&run events) and two lasting single-frame fusion events. The trajectory is convoluted with an idealized step function g(t), which is run over the entire time series - here shown for snapshots at time t=5 and t=160. The signal convolution will result in a trajectory of the intersection between the two functions (f(t)∗g(t)) resulting in a spike at the position of the docking event, due to the maximum overlap between the two functions. On top of the filtered signal convolution trajectory, a third power function is applied to enhance the signal and eliminate noise followed by peaks detected using the SciPy signal processing package. The positions for the peaks are saved and used to attain the raw intensities at the original trajectories for further analysis and classification.

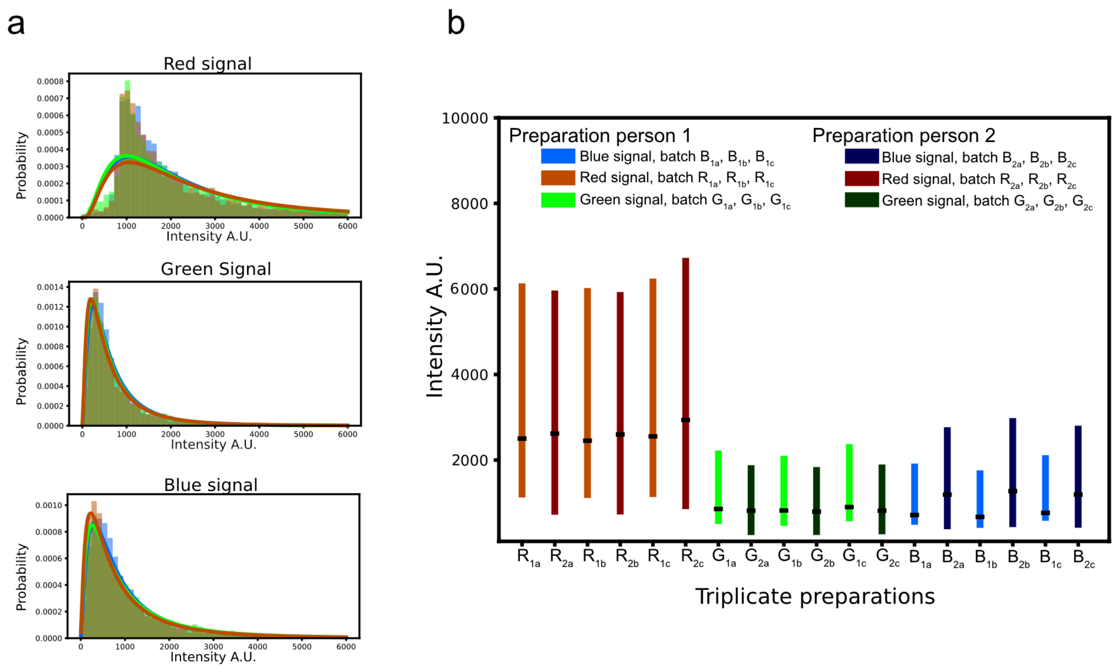


#### Supplementary Figure 11 – Reproducibility in preparation of liposomes, 3 biological repetitions and imaging by two independent persons

Reproducible liposome preparation is essential for the method, so as to use the liposome for training a machine learning model and recognize the fluorescent lipid composition. Two individual persons prepared and imaged independently three 3 biological repetitions of the liposome chromophore composition referred to as (5,5,5) as it contains three different fluorescent lipids and therefore potentially the most heterogenic liposome composition. (a) The intensity profile for all three populations is plotted and individual fitted using a lognormal distribution in all three imaging channels (Red, Green, Blue). (b) Mean and standard deviation for each fit for each channel are plotted for both test persons show casting robust and reproducible preparation, and imaging.

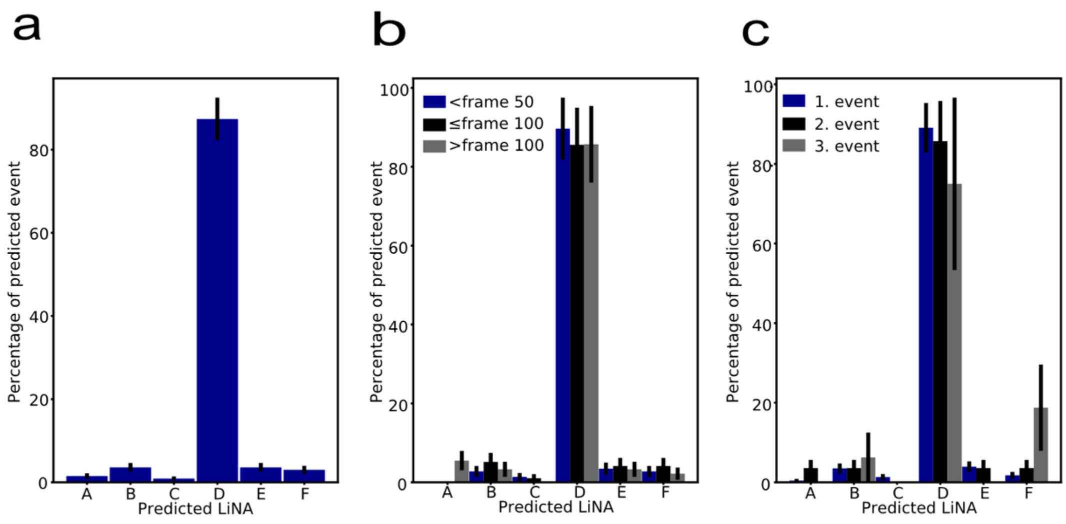


Supplementary Figure 12 – Ground truth data validation of the machine learning model performance with liposomes associated with LiNA D' and fluorescently barcoded (5,5,5).

a) predictions for liposomes barcoded with (5,5,5) and LiNA D'. The ML model correctly predicts the liposomes in 92.3 % of cases, (see confusion matrix in Fig. 2a). On dynamic data, we find 87.4±5.1 % (Supplementary Fig. 11a), which within error fits with the expected from the model. 3.6±1.0 % of the data are classified as B' or E', and 3.0±0.9 % as F, which further both follows and verifies the model on dynamic data. b), prediction accuracy is found to be independent of whether fusion occurs in the first or last part of the experiments, by dividing the event into the occurrence time with equal amount of data in each. c) the prediction accuracy is independent of the order of events. The larger uncertainty for the third event is due to less data than for the first and second event.

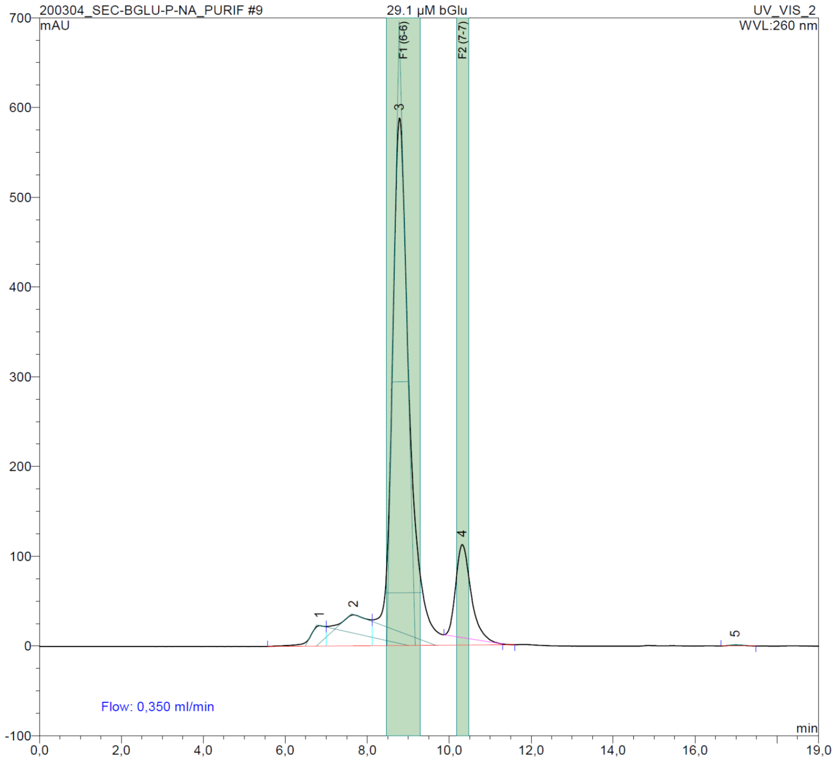


#### Supplementary Figure 13 – Purification of β-glucosidase

Size-exclusion HPLC chromatogram of 60 µl β-glucosidase concentrated stock solution (29 µM). Peaks 1 + 2 were not collected. Only Peak 3 exhibited enzyme activity (see methods) and its retention time matches well with the molecular mass of the enzyme dimer (121 kDa). Peak 4 stems from bovine serum albumin (67 kDa, stabilizer additive from the supplier).

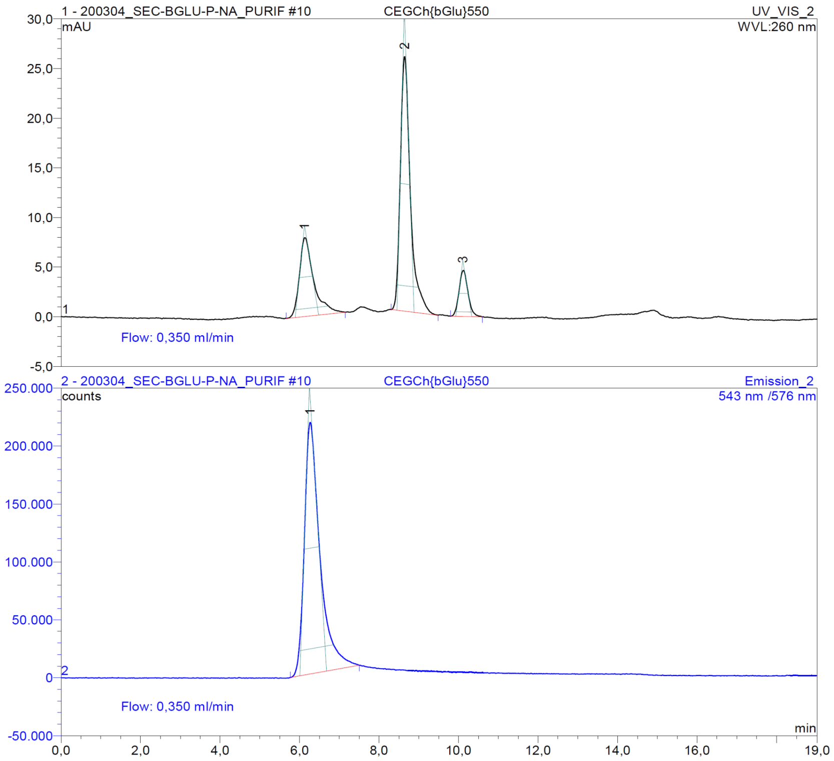


#### Supplementary Figure 14 – Example size exclusion of liposomes encapsulating βGlu

ATTO-550-labeled target liposomes rehydrated in presence of βGlu, injected after extrusion to 100 nm (figure shows analytic run, overlay of UV and fluorescence traces). Peak 1 (5.8 - 6.4 min, identified via the ATTO-550 fluorescence (Ex. 543 nm, Em. 576 nm), bottom panel) contained liposomes and was collected for fusion experiments in bulk.

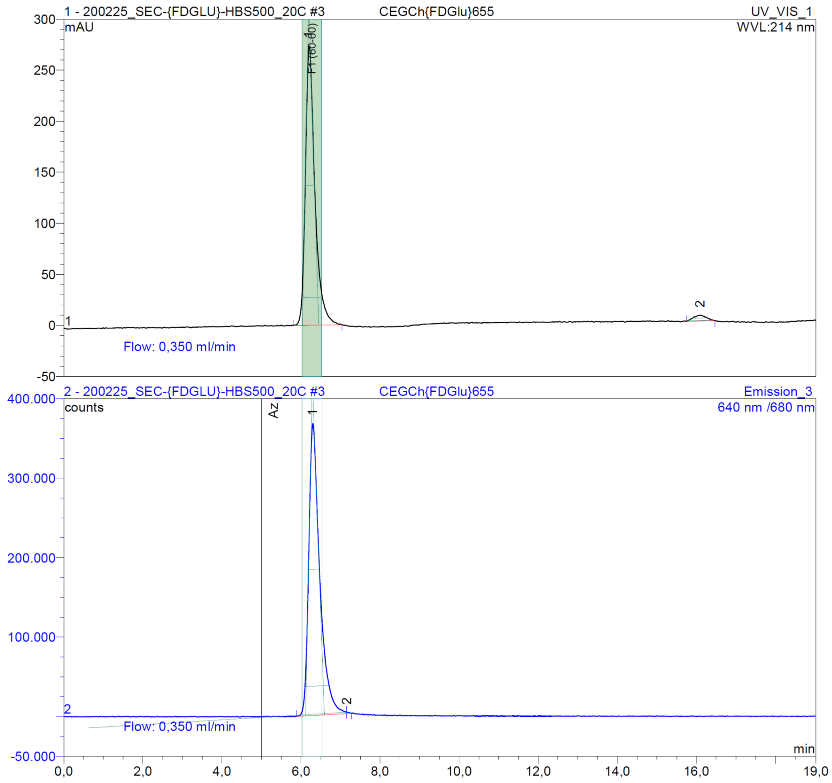


#### Supplementary Figure 15 – Example size exclusion of liposomes encapsulating FDGlu

ATTO-655-labeled target liposomes rehydrated in presence of βGlu, and injected after extrusion to 100 nm (figure shows analytic run, overlay of UV absorbance and fluorescence traces). Peak 1 (6.0-6.8 min, identified via the ATTO-655 fluorescence (Ex. 640 nm, Em. 680 nm), bottom panel) contained liposomes and was collected for leakage and fusion experiments in bulk. Peak 2 at 214 nm shows unentrapped FDGlu (which has a very low extinction coefficient).

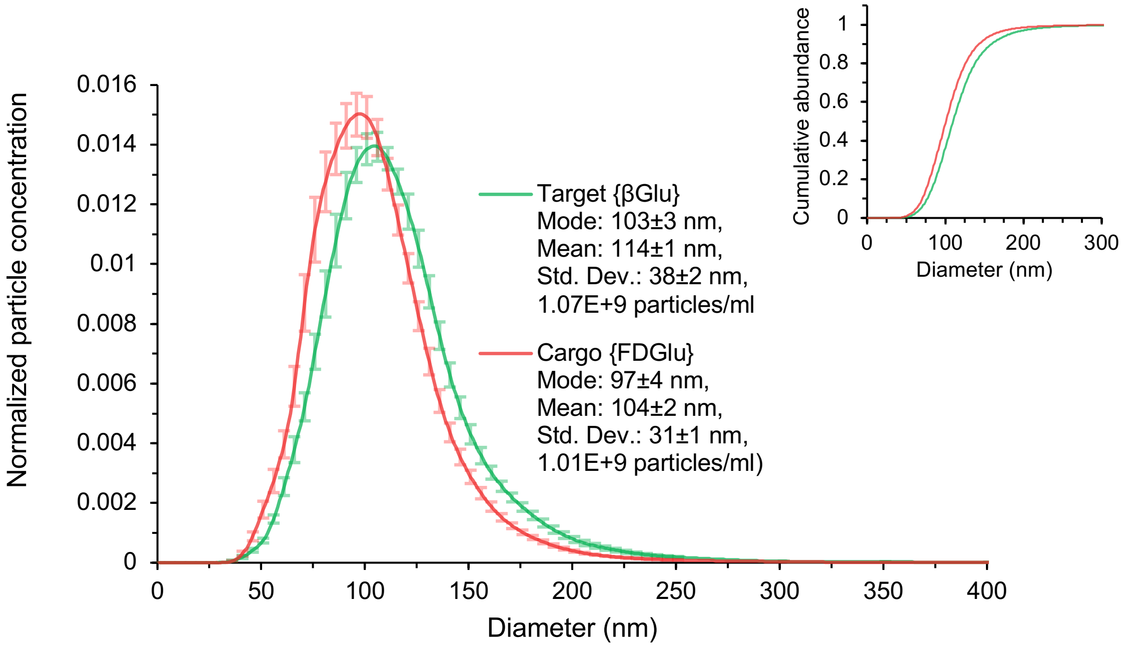


#### Supplementary Figure 16 – Liposome characterization using Nanoparticle Tracking Analysis (NTA)

Size distribution data for liposomes used for the TIRF/bulk fusion experiments, obtained after size-exclusion chromatography: liposomes encapsulating β-Glucosidase ({βGlu}, labeled with 0.1 mol% biotin-PEG2000-DOPE, 0.5 mol% ATTO-550-DOPE) or the substrate fluorescein-di-glucopyranoside ({FDGlu}, labeled with 0.5 mol% ATTO-655-DOPE). The plot shows the envelope of the numerical size distribution at 1 nm intervals (5 nm moving average) based on the average (± std. err.) of 10 measurements. The numerical standard deviation of the distribution (38 nm and 31 nm for {βGlu} and {FDGlu}, respectively) indicates a low polydispersity, albeit with a small sub-population below 60 nm (2.2% for {βGlu} and 3.8% for {FDGlu}, and above 200 nm (2.8% for {βGlu} and 1.1% for {FDGlu}, see inset with cumulative distribution. The Brownian motion of individual liposomes was recorded using a NanoSight LM10-HS microscope equipped with an Andor Lucas EMCCD camera, a LM14 temperature controller. Liposomes were made visible via their scattering of a shallow angle laser beam (diode laser, 404 nm). The HPLC fractions from SEC were diluted with using freshly filtered (0.1 µm) HBS to a concentration of approx. $10^9$ particles/ml (850-950 µM to 2 µM lipid). More than 5000 individual particle diffusion tracks were recorded (20 °C, 25 fps, camera gain 300, 10 videos of 15 s). The data was analyzed using the NanoSight NTA 2.3 software (detection threshold 6, min. track length 8, max. jump distance <12.5 nm).

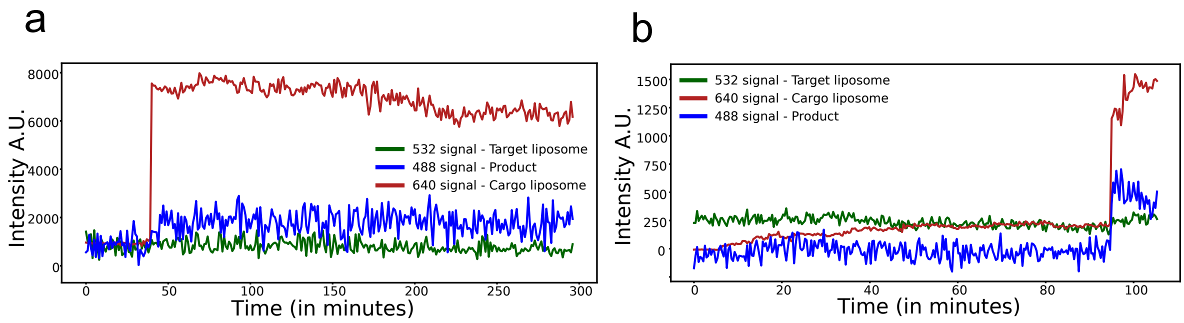


#### Supplementary Figure 17 – Representative Trajectories of successful cargo delivery by DNA mediated fusion

Cargo liposomes containing LiNA complementary to the functionalization on the target liposomes are allowed to fuse with Target liposomes are loaded with β-glucosidase and membrane labelled using ATTO-550-DOPE for localization. Cargo liposomes are membrane labelled using ATTO-655-DOPE and loaded with a pro-fluorescent substrate, FDGlu. Docking of the cargos will thereby result in a single step increase in the red channel. The successful fusion will result in cargo delivery and enzymatic reaction resulting in an increase in the blue channel. The two trajectories are random samples from successful fusion, showing increase in blue channel quickly after liposome docking (red).

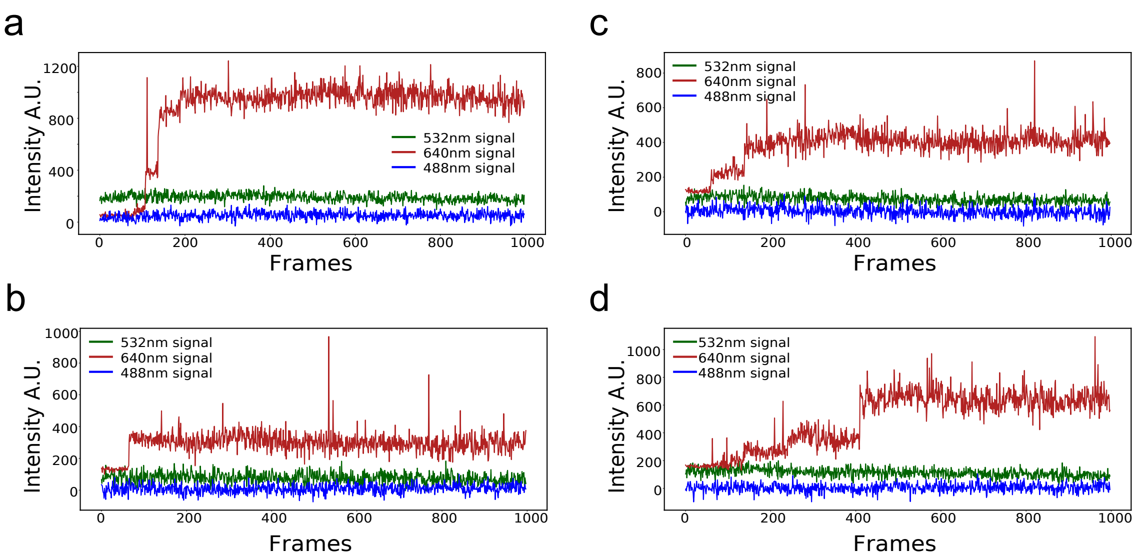


Supplementary Figure 18 – Representative Trajectories of control experiments displaying liposome fusion for empty cargo liposomes with complimentary LiNA sequences showing no product formation

Target liposomes and cargo liposomes are prepared as described in both method with targets loaded with β-glucosidase and membrane labelled using ATTO-550-DOPE and with cargo membrane labelled using ATTO-655-DOPE but without any loading of pro-fluorescent substrate. Upon docking of cargo as detected in the red channel no conversion of substrate occurs, allowing us to correct for fluorescent emission cross talk. From the trajectories we see multiple single-step docking, and no observed changes in the blue product channel. We find 89.96±6.14% docking, which within error bars are in agreement with cargo vesicles loaded with pro-fluorescent substrate. The 4.0 ± 1.4% as a false-positive product conversion, giving the low cross talk of the method.

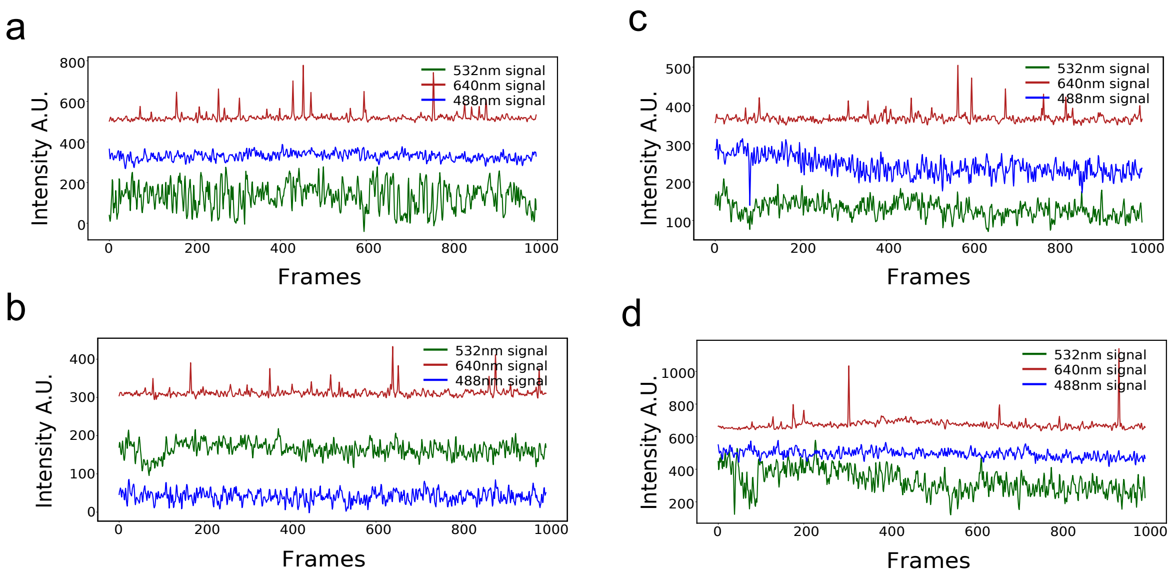


#### Supplementary Figure 19 – Representative Trajectories of control experiments of cargo liposomes with non-complementary LiNA sequences displaying no prolonged docking and thus no fusion events

Target liposomes and cargo liposomes are prepared as described in both methods, with targets loaded with β-glucosidase and membrane labelled using ATTO-550-DOPE and cargo membrane labelled using ATTO-655-DOPE and loaded with a pro-fluorescent substrate, FDGlu. Here cargo liposomes are functionalized with a LiNA sequence non-complementary to the target sequence. As seen from the traces, cargo vesicles do kiss and run on the target liposomes giving rise to all the spikes observed in the red channel. 1.9$\pm$0.8 % of the cargo liposomes docks, but 0% leads to fusion detected in the blue product channel.

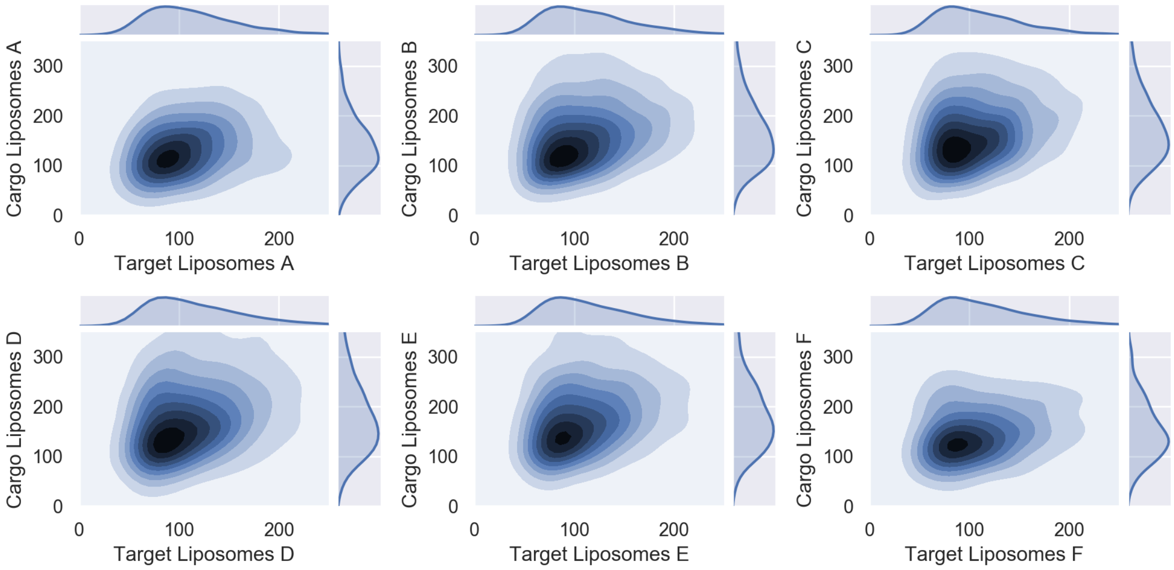


#### Supplementary Figure 20 – Relation between cargo liposome size and target liposome size

3D plots displaying the relation between the liposome's sizes of target and cargo liposomes. Pseudocolor corresponds to probability density. A slightly positive correlation with a Pearson correlation of 0.2 for all six populations, but with broad distributions. This shows that all targeted liposomes can facilitate fusions for all cargo liposomes across sizes with a small preference for large targets to fuse with large cargos.

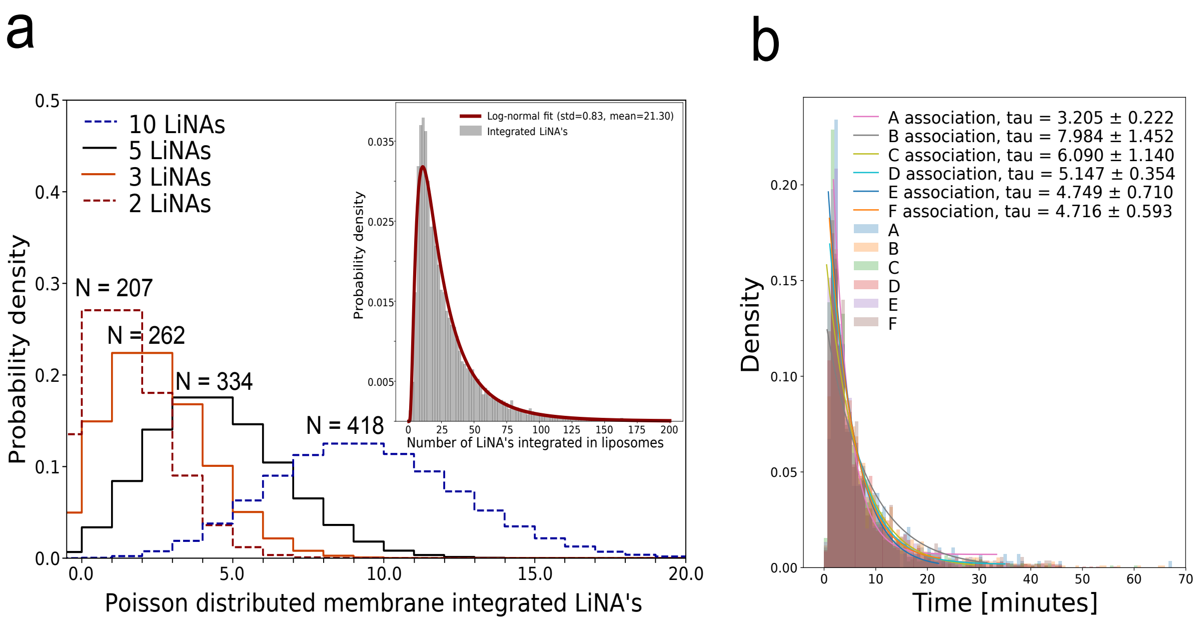


#### Supplementary Figure 21– Quantification of Docking lag time and fusion for individual LiNA sequences

a) The individual number of LiNA sequences per liposome can be calculated assuming an equal incorporation of LiNA decoration across different liposome sizes. Assuming incorporation follows a Poisson distribution and analyzing all fusion events, we find 207 cargo liposomes have fused with only one available LiNA. 262 cargo fused with two LiNA present, 334 with tree LiNAs and 418 cargos with four liNAs present. This suggest and supports that only a few single digit LiNAs are sufficient for facilitation of fusion[6]. b) The automated ML classification allowed us to deconvolute fusion kinetics for the different LiNA sequences. The distribution of the docking lag of cargos, followed an exponential decay, which confirmed that docking occurs in a single step process. Interestingly, docking mediated by A-A’ hybridization was found to be the fastest with a waiting time of 3.2±0.2 minutes after cargo liposome addition, while the B-B’ pair was the slowest with an average waiting time of 8.0±1.4 minutes for identical concentrations. Previously it was shown that the waiting time depended on the number of DNA strands participating in the fusion process [6,7], which is a good agreement with our finding.

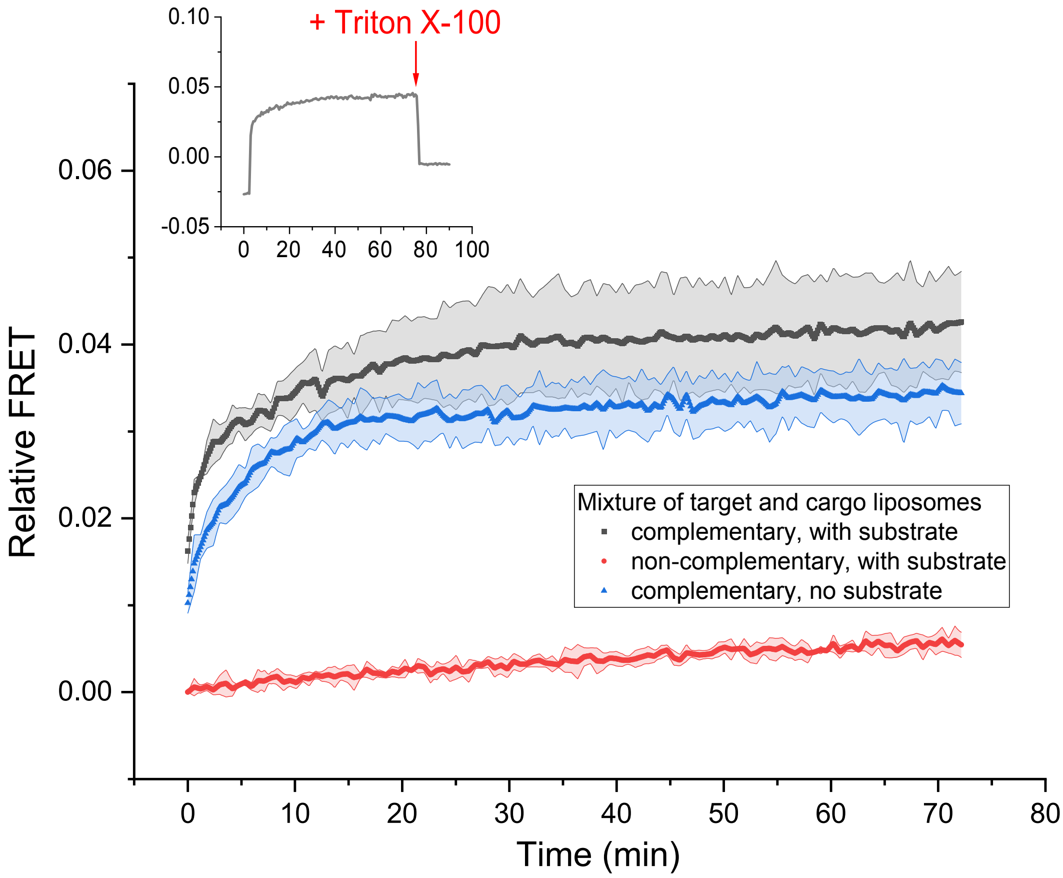


#### Supplementary Figure 22 - Lipid mixing monitored by FRET in spectrometer

Lipid mixing monitored by Förster Resonance Energy Transfer (FRET) between the lipid dyes of the liposomes that contain enzyme and substrate. Upon fusion, lipids of the Atto-550-DOPE labeled βGlu loaded liposomes and the Atto-655-DOPE labeled substrate or empty liposomes serving as control for crosstalk on FRET upon substrate mixing). Only in case of having complementary LiNAs (D and D’) on the surface of the population, the intensity in the accepter channel (excitation (Ex.) 532 nm, emission (Em.)680 nm) increased significantly. The result underlines the kinetic observation during microscopy, that most of the possible fusion events are done within 30 min. The data were offset-corrected to the $I_0$ of the non-complementary control. Addition of 0.1 % w/v Triton-X 100 completely disrupted the FRET signal (inset). The relative FRET efficiency was calculated as $E = (A-A_0)/(D+(A-A_0))$, where A is the acceptor fluorescence intensity (532/680 nm), $A_0$ the acceptor intensity (532/680 nm) at $t_0$ for the non-complementary experiment (baseline correction) and D is the donor intensity (532/576 nm).

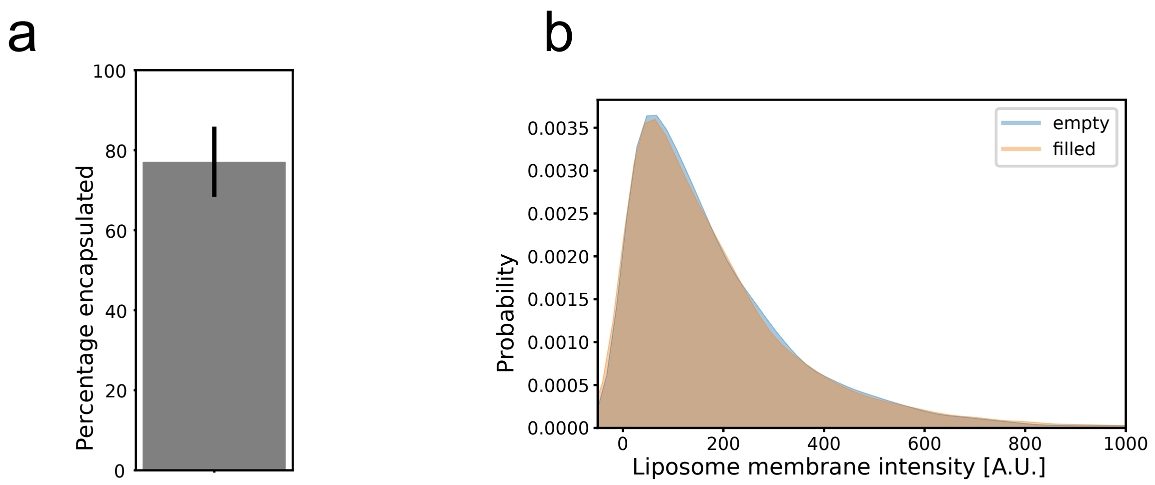


#### Supplementary Figure 23 – Encapsulation efficiency of β-glucosidase

a) Encapsulation efficiency of fluorescently labelled β-glucosidase to labeled liposomes (see Methods for preparation). Analysis of a total of 11791 liposomes, reveals an encapsulation efficiency of 77.13%. b) Liposome sizes distributions are identical for liposomes filled with β-glucosidase and empty liposomes. Comparing the background encapsulation channel for both filled and empty liposomes we can calculate the percentage of liposomes with encapsulated β-glucosidase. Analyzing 11083 liposomes, 5178 filled and 5905 empty liposomes, we find an encapsulation efficiency of 77.13% as shown in (a).

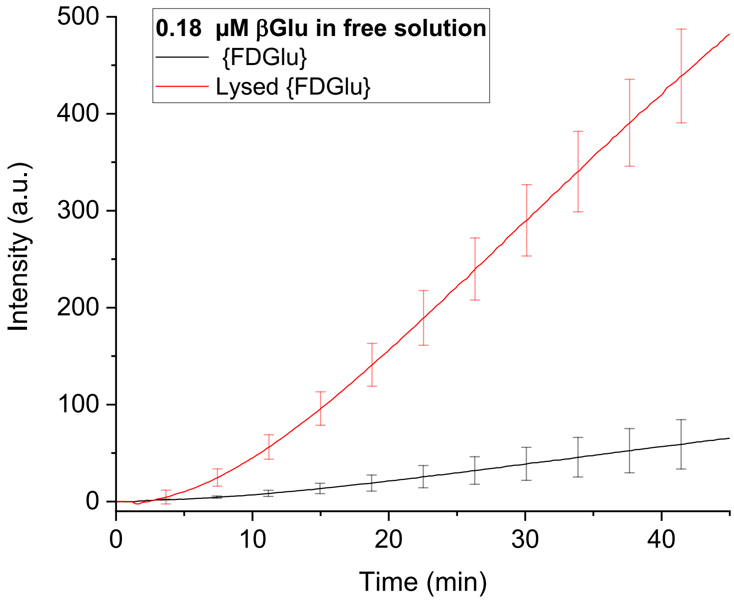


#### Supplementary Figure 24 – Leakage of FDGlu substrate recorded by spectrometer

100 µM lipid encapsulating FDGlu was incubated at 37 °C in presence of a high concentration of freely dispersed βGlu (~100 fold more than in fusion experiments). Fluorescence intensity monitored for the fluorescein product (Ex. 488 nm and Em. 510 nm) reveals minor leaking in the spectrometer assay, as compared to liposomes lysed by detergent. Error bars = standard deviation of 3 independent measurements

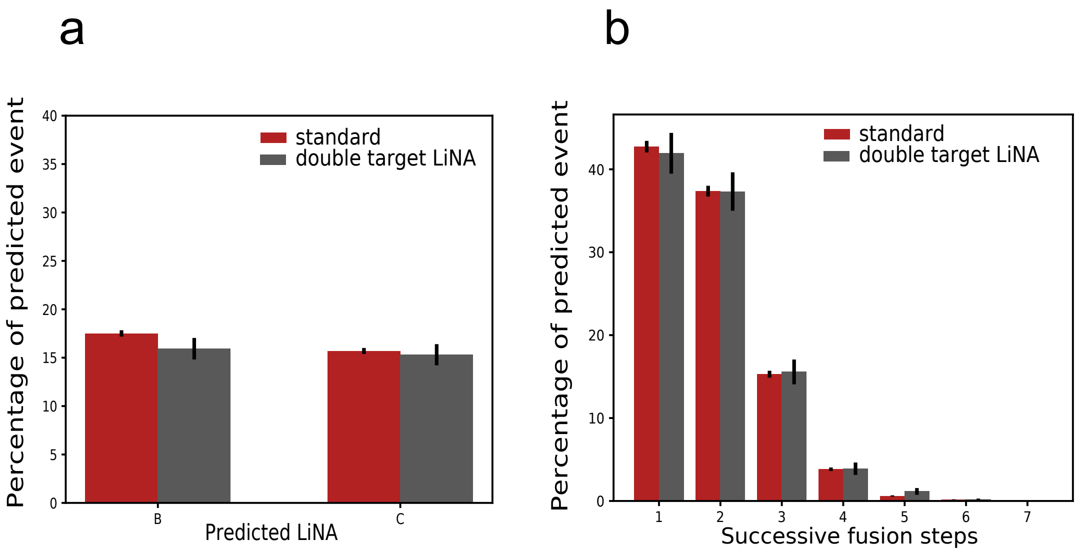


Supplementary Figure 25 – Number of docking and fusion events are not limited by LiNA depletion

a) Doubling the amount of LiNA functionalization (gray barplots) on cargo vesicles, does not change the occurrence profile (compared to the red barplot). Data shown for B and C LiNA functionalized liposomes. b) Doubling the amount of the six LiNA sequences on all targets (gray barplot, compared to red barplot) does not alter measurably the number of successive fusion events. Data for 8830 target liposomes which was subjected to 16143 individual fusion of cargos

#### Supplementary Table 1 – Possible distinct permutations for different sizes of cargo libraries for multiplexing assay

| N<br># Subsequent events | $3^N$<br># Possible distinct permutations | $4^N$<br># Possible distinct permutations | $5^N$<br># Possible distinct permutations | $6^N$<br># Possible distinct permutations |
|---|---|---|---|---|
| 1 | 3 | 4 | 5 | 6 |
| 2 | 9 | 16 | 25 | 36 |
| 3 | 27 | 64 | 125 | 216 |
| 4 | 81 | 256 | 625 | 1296 |
| 5 | 243 | 1024 | 3125 | 7776 |
| 6 | 729 | 4096 | 15625 | 46656 |
| 7 | 2187 | 16384 | 78125 | 279936 |

Maximum number of possible distinct permutations for cargo fusions follows a Power law dependence on both number of LiNA sequences (ɣ) and number of subsequent fusion events (N). This allows recording of $ɣ^N$ distinct combinatorial fusions. We have shown the robustness of the method for six barcodes and six associated distinct LiNA sequence. The table summarizes the number of possible distinct permutations for three to 6 six barcodes (and LiNAs) with one to 7 subsequent fusion events. From the table it can be seen that both number of barcodes ɣ and subsequent events (N) are important for establishing a high throughput method.
